# Autolysosomal exocytosis of lipids protect neurons from ferroptosis

**DOI:** 10.1101/2022.11.24.517842

**Authors:** Isha Ralhan, Jinlan Chang, Matthew J. Moulton, Lindsey D. Goodman, Nathanael Y.J. Lee, Greg Plummer, H. Amalia Pasolli, Doreen Matthies, Hugo J. Bellen, Maria S. Ioannou

**Affiliations:** Department of Physiology, University of Alberta, Edmonton, AB T6G 2R3, Canada; Group on Molecular and Cell Biology of Lipids, University of Alberta, Edmonton, AB T6G 2R3, Canada; Department of Molecular and Human Genetics, Baylor College of Medicine, Houston, TX 77030, USA; Jan and Dan Duncan Neurological Research Institute, Texas Children’s Hospital, Houston, TX 77030, USA; Faculty of Medicine & Dentistry Cell Imaging Core, University of Alberta, Edmonton, AB T6G 2R3, Canada; Electron Microscopy Resource Center, The Rockefeller University, New York, NY 10065, USA; Janelia Research Campus, Howard Hughes Medical Institute, Ashburn VA 20147, USA; Unit on Structural Biology, Division of Basic and Translational Biophysics, Eunice Kennedy Shriver National Institute of Child Health and Human Development, National Institutes of Health, Bethesda MD 20847, USA; Department of Neuroscience, Baylor College of Medicine, Houston, TX 7703, USA; Department of Cell Biology, University of Alberta, Edmonton, AB T6G 2R3, Canada; Neuroscience and Mental Health Institute, University of Alberta, Edmonton, AB T6G 2R3, Canada

## Abstract

During oxidative stress neurons release lipids that are internalized by glia. Defects in this coordinated process play an important role in several neurodegenerative diseases. Yet, the mechanisms of lipid release and its consequences on neuronal health are unclear. Here, we demonstrate that lipid-protein particle release by autolysosome exocytosis protects neurons from ferroptosis, a form of cell death driven by lipid peroxidation. We show that during oxidative stress, peroxidated lipids and iron are released from neurons by autolysosomal exocytosis which requires the exocytic machinery; VAMP7 and syntaxin 4. We observe membrane-bound lipid-protein particles by TEM and demonstrate that these particles are released from neurons using cryoEM. Failure to release these lipid-protein particles causes lipid hydroperoxide and iron accumulation and sensitizes neurons to ferroptosis. Our results reveal how neurons protect themselves from peroxidated lipids. Given the number of brain pathologies that involve ferroptosis, defects in this pathway likely play a key role in the pathophysiology of neurodegenerative disease.

**SUMMARY:** Release of lipid-protein particles by autolysosomal exocytosis protects neurons from ferroptosis.

## INTRODUCTION

Oxidative stress and the generation of high levels of reactive oxygen species (ROS) have been associated with nearly all known neurodegenerative diseases. The damaging effects of ROS on proteins and nucleic acids in the brain are well characterized. The effects of ROS on lipid homeostasis have recently gained attention as a critical feature in neurological disease. Yet many of the mechanisms that cells use to protect themselves from damaged lipids during oxidative stress are unknown.

Excessive levels of ROS attack lipids in a process called lipid peroxidation. Polyunsaturated fatty acids on membrane phospholipids are the preferred lipid substrate for ROS (Kagan et al., 2017). The resulting lipid peroxides, if not neutralized by the cell’s antioxidant defense system, can induce a biochemically distinct form of cell death called ferroptosis (Dixon et al., 2012). The brain is enriched with polyunsaturated fatty acids that are rapidly esterified to membrane phospholipids (Bazinet & Layé, 2014) and neurons constantly generate and utilize ROS as a by-product of metabolism and as an important signaling molecule (Cw Oswald et al., 2018) This makes neurons highly susceptible to lipid peroxidation and consequently, ferroptosis (Agmon et al., 2018; Yang & Stockwell, 2016)

During oxidative stress, neurons use autophagy to remove portions of organelles and/or membranes including those containing peroxidated lipids (Ioannou, Jackson, et al., 2019). Lysosomal degradation of membranes during autophagy is often accompanied by an increase in fatty acid storage as neutral lipids within lipid droplets (Rambold et al., 2015). Neurons, however, make only a limited number of lipid droplets (Ralhan et al., 2021). Instead, neurons transport excess fatty acids to glial cells (Ioannou, Jackson, et al., 2019; Liu et al., 2015, 2017). Glia internalize fatty acids by endocytosis and store them in lipid droplets (Ioannou, Jackson, et al., 2019; Moulton et al., 2021). This process plays a clear role in protecting the brain as failure to form lipid droplets in glia results in neurodegeneration (Liu et al., 2017; Moulton et al., 2021). Despite the importance of these processes, the mechanisms of fatty acid release from neurons are poorly characterized.

One way that neurons release fatty acids is through adenosine triphosphate-binding cassette transporter A1 (*ABCA1*) and A7 (*ABCA7*) (Moulton et al., 2021). In order for fatty acids generated by autophagy to be released from neurons via plasma membrane transporters, they would need to exit lysosomes and related compartments. During cellular stress, however, lysosomes, late endosomes, and/or autophagic vesicles can fuse directly with the plasma membrane (Buratta et al., 2020; Hedlund et al., 2011; Ravi et al., 2016) Autolysosome exocytosis is becoming increasingly recognized for its effects on cellular health, particularly in the context of lysosomal storage disorders (Spampanato et al., 2013) where undigested and peroxidated lipids accumulate together with proteins in autolysosomal compartments. This raises the possibility of autolysosomal exocytosis as an alternate mechanism for neuronal lipid release.

Here, we investigate novel mechanisms and function of neuronal lipid release during cellular stress. We demonstrate that neuronal release of peroxidated lipids and iron occurs by autolysosomal exocytosis and is dependent on the SNARE complex proteins VAMP7 and syntaxin 4. Neurons contain lipid-protein particles within autolysosomes and these lipid-protein particles are released by neurons. Finally, release of lipid-protein particles by exocytosis protects neurons from accumulation of peroxidated lipids, iron, and ferroptosis. Together, these findings reveal autolysosomal exocytosis as a critical mechanism to protect neurons from peroxidated lipids during oxidative stress.

## RESULTS

### Neurons release fatty acids by exocytosis

We explored whether exocytosis contributes to the release of fatty acids from neurons during oxidative stress. Assays were performed with hippocampal neurons grown in cytosine arabinoside to eliminate proliferating glia thereby generating cultures where 95% of cells stain positive for neuron-specific β-tubulin-III (Fig. S1, A and B). Neurons release lipids in Hanks’ Balanced Salt Solution (HBSS) (Ioannou, Jackson, et al., 2019), which, in addition to starvation, induces oxidative stress, in the form of superoxide and hydroxyl radicals (Choi et al., 2015), as indicated by increased CellRox staining (Fig. S1, C and D). As exocytosis is a calcium-regulated process, we tested whether modulating calcium levels affect fatty acid release from neurons in HBSS. We loaded neurons with BODIPY 558/568 C12 (Red-C12), a fluorescently labelled, saturated fatty acid, and treated neurons with HBSS plus BAPTA-AM, a cell permeable calcium chelator that inhibits exocytosis (Pryor et al., 2000). We found that inhibiting exocytosis with BAPTA-AM increased Red-C12 accumulation within neurons revealed by confocal microscopy (Fig. 1, A and B and S1, E) and decreased Red-C12 released into the conditioned-HBSS as detected using a fluorescence plate-reader (Fig. 1 C). We also treated neurons in HBSS plus MK6-83 which stimulates exocytosis (Tsunemi et al., 2019). Specifically, MK6-83 activates transient receptor potential mucolipin cation channels on endosomal membranes, mediating calcium release for exocytosis. Stimulating exocytosis with MK6-83 resulted in a decrease in neuronal Red-C12 accumulation (Fig. 1, A and B) and an increase in Red-C12 released (Fig. 1 D). As fatty acids released by neurons during oxidative stress are internalized by neighbouring glia (Ioannou, Jackson, et al., 2019), we next examined the effects of MK6-83 on fatty acid transport to glia. We loaded neurons with Red-C12, cultured them with unlabelled glia on a separate coverslip separated by paraffin wax, and assessed the appearance of Red-C12 puncta within glia (Fig. 1 E). Consistent, with an increase in Red-C12 release, MK6-83 increased neuron-derived Red-C12 detected in glia (Fig. 1, F and G).

**Figure 1.**
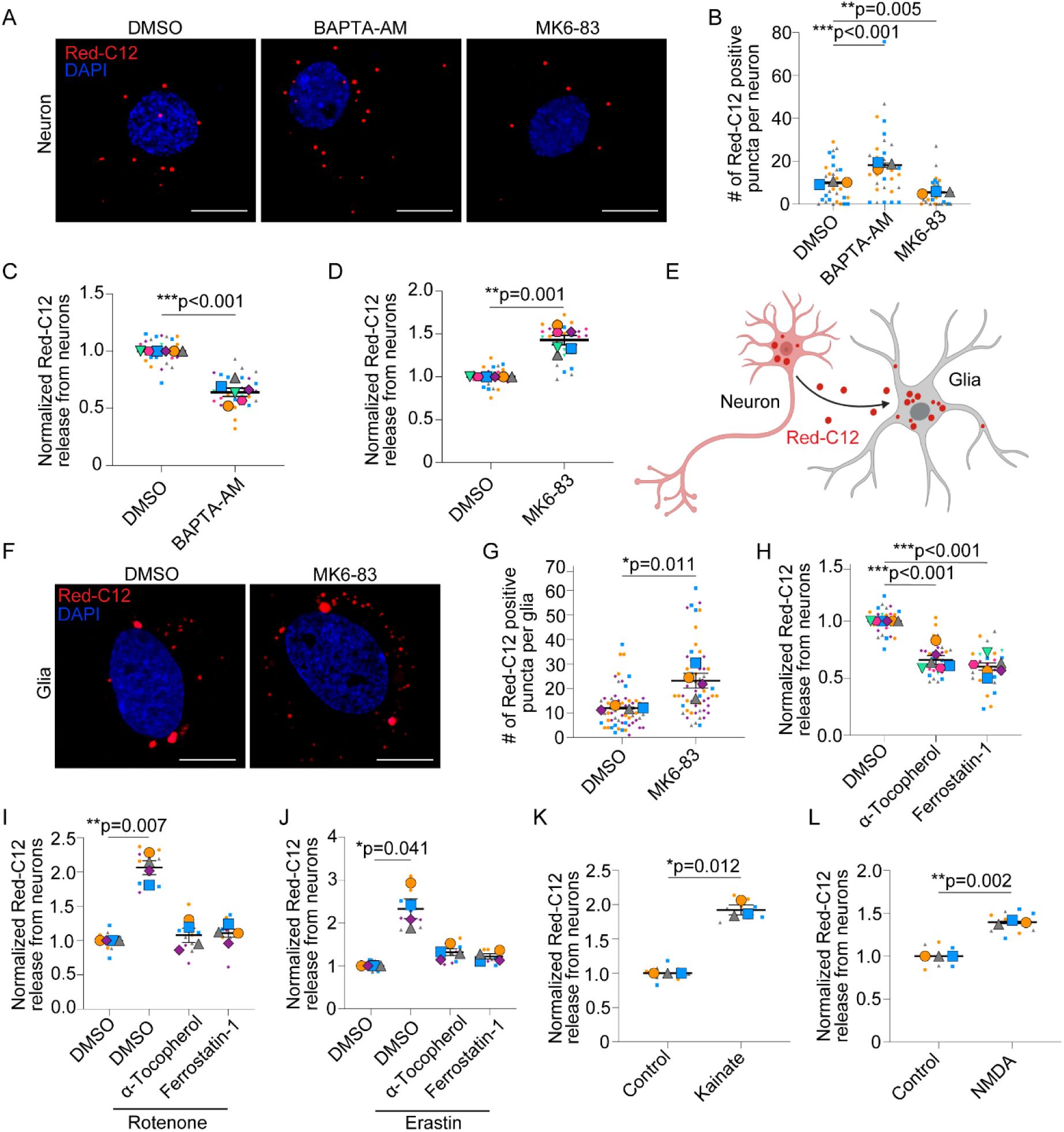
Neurons release fatty acids by exocytosis. (A) Confocal maximum intensity projection of neurons loaded with Red-C12 and treated with HBSS ± DMSO, BAPTA-AM or MK6-83. Scale bars, 10 ∝m. (B) Red-C12 positive puncta per neuron. n = 3 independent experiments; mean ± SEM. One-way ANOVA with Dunnett’s post test. (C) Neuron-conditioned HBSS analyzed for Red-C12 fluorescence and normalized to DMSO treated neurons. n = 6 independent experiments; mean ± SEM. One sample t-test with Bonferroni correction. (D) Neuron-conditioned HBSS analyzed for Red-C12 fluorescence and normalized to DMSO treated neurons. n = 6 independent experiments; mean ± SEM. One sample t-test adjusted for multiple comparisons using Bonferroni correction. (E) Schematic depicting Red-C12 transfer assay from neurons to glia. (F) Confocal image of glia post-transfer assay in HBSS ± DMSO or MK6-83. Scale bars, 10 ∝m. (G) Red-C12 positive lipid droplets in glia following fatty acid transfer assay. n = 4 independent experiments; mean ± SEM. Student’s t-test. (H) Neuron-conditioned HBSS analyzed for Red-C12 fluorescence and normalized to DMSO treated neurons. n = 6 independent experiments; mean ± SEM. One sample t-test with Bonferroni correction. (I-J) Neuron-conditioned media analyzed for Red-C12 fluorescence and normalized to DMSO treated neurons. n = 4 independent experiments; mean ± SEM. One sample t-test with Bonferroni correction. (K-L) Neuron-conditioned media analyzed for Red-C12 fluorescence and normalized to control neurons. n = 3 independent experiments; mean ± SEM. One sample t-test.

Treating with α-tocopherol or ferrostatin-1, lipophilic antioxidants, decreased Red-C12 released from neurons in HBSS, suggesting the importance of oxidative stress for exocytosis (Fig. 1 H). To tease apart the contribution of oxidative stress from starvation, we measured Red-C12 release from neurons upon ROS induction in complete media. Red-C12 release increased following treatment with rotenone, a mitochondria complex I inhibitor, and erastin, a ferroptosis activator, which could similarly be rescued with α-tocopherol and ferrostatin-1 (Fig. 1, I and J). Similarly, Red-C12 release increased with excitotoxicity induced with kainate or N-methyl-D-aspartate (NMDA) (Fig. 1, K and L).

We then used total internal reflection fluorescence (TIRF) microscopy to visualize Red-C12 exocytic events. We detected several Red-C12 puncta at the plasma membrane compatible with exocytosis following treatment with MK6-83 (Fig. 2, A and B). These fusion events exhibit similar kinetics to exocytic events previously described in cultured neurons (Fig. 2 C) (Roman-Vendrell et al., 2014). Neurons treated with MK6-83 to stimulate exocytosis had more Red-C12 puncta at the plasma membrane by TIRF microscopy than those treated with BAPTA-AM, which inhibits exocytosis (Fig. 2 D). Collectively, these results support exocytosis as an important mechanism for fatty acid release from neurons.

**Figure 2.**
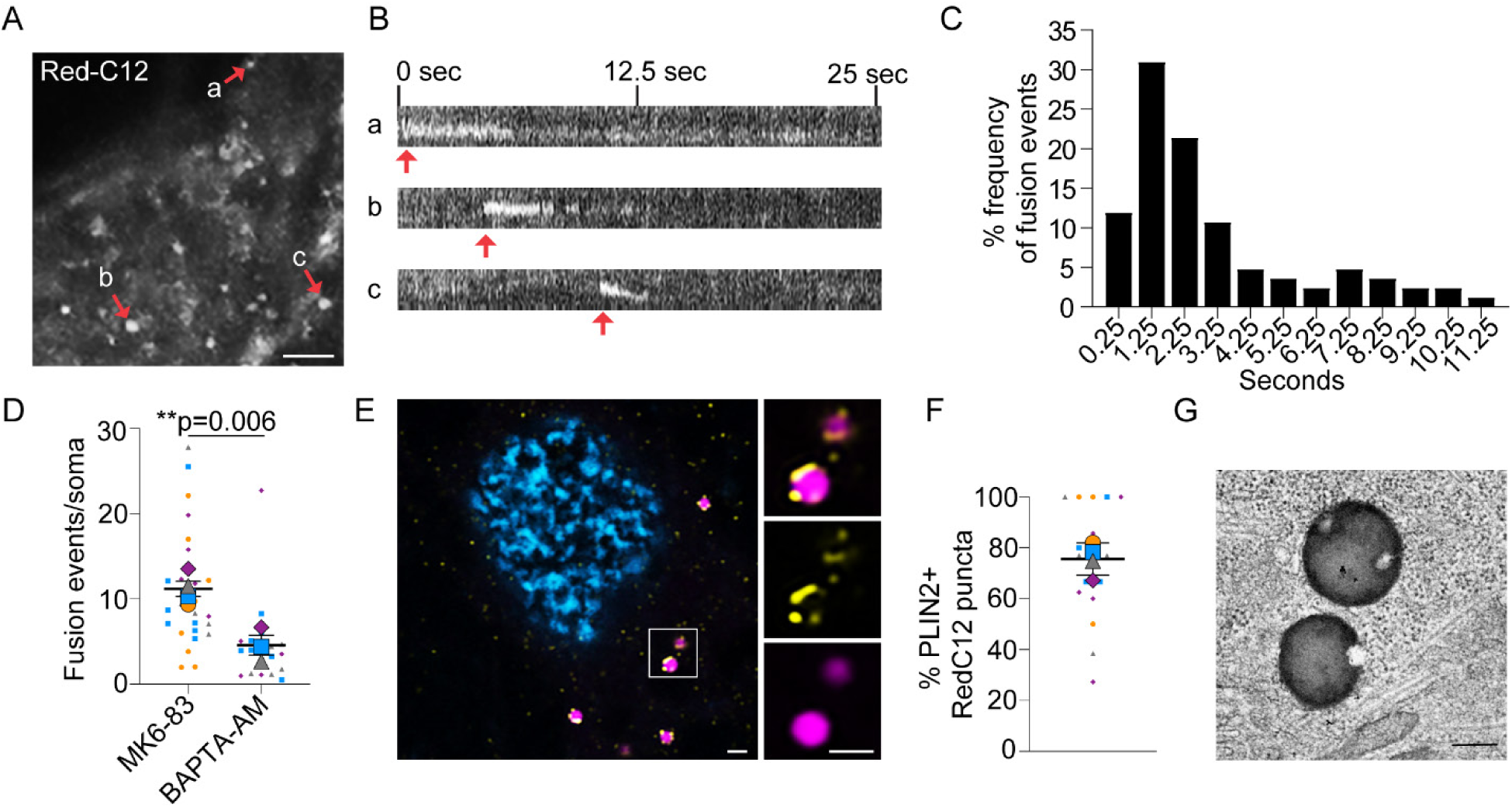
Neurons exocytose Red-C12 by TIRF microscopy. (A and B) Live neurons in Tyrode’s solution containing MK6-83 imaged by TIRF microscopy. (A) Maximum intensity projection over a 54 sec interval. Scale bar, 2.5 ∝m. (B) Kymograph, red arrows show select exocytic events. (C) Frequency of exocytic events in neurons + MK6-83. n = 84 events from 10 cells in 2 independent experiments. (D) Neurons ± MK6-83 or BAPTA-AM in HBSS were analyzed for exocytic events using TIRF microscopy. The number of fusion events per cell was normalized to soma size. n = 3-4 independent experiments; mean ± SEM. Student’s t-test. (E) Airyscan image of neurons in HBSS labelled with Red-C12 and immunostained for PLIN2. Boxed area magnified in right panels. Scale bars, 1 ∝m. (F) Percent colocalization of PLIN2 and Red-C12 positive puncta in (E). n = 4 independent experiments; mean ± SD. (G) Lipid droplets in neurons stained with imidazole-buffered osmium and imaged by transmission electron microscopy. Scale bar, 250 nm.

We next identified several of the Red-C12 puncta within neurons as lipid droplets by colocalization of Red-C12 with the lipid droplet marker PLIN2 (Fig. 2, E and F). The presence of bona fide lipid droplets in cultured neurons was further confirmed by staining unsaturated fatty acids with imidazole-buffered osmium and imaging by transmission electron microscopy (Fig. 2 G). While lipid droplets rarely form in neurons in vivo (Ralhan et al., 2021), their formation in vitro serves as an important readout for fatty acid accumulation. This suggests that fatty acids that fail to be released can be routed to and/or remain in lipid droplets for storage in cultured neurons.

### Neurons release fatty acids by lysosomal exocytosis

As neurons release pathogenic proteins via lysosomal exocytosis (Tsunemi et al., 2019), we explored whether lysosomal exocytosis similarly participates in neuronal fatty acid release. To do this, we knocked down the soluble N-ethylmaleimide sensitive factor attachment protein receptor (SNARE) complex required for lysosomal fusion with the plasma membrane; vesicle-associated membrane protein 7 (VAMP7) on lysosomes and syntaxin 4 on the plasma membrane (Rao et al., 2004). Using lentivirus to deliver three independent shRNAmiRs for each target protein, we reduced expression of VAMP7 or syntaxin 4 compared to a nontargeting shRNAmiR control (Fig. 3, A-D). We then loaded neurons with Red-C12 overnight and examined the amount of Red-C12 released. In the absence of VAMP7 or syntaxin 4, Red-C12 release in HBSS was reduced (Fig. 3, E and F). Syntaxin 4 knockdown similarly prevented Red-C12 release into complete media upon treatment with oxidative stressors rotenone, erastin and kainate (Fig. 3, G-I). Syntaxin 4 knockdown also decreased Red-C12 at the plasma membrane as detected by TIRF microscopy in the presence of MK6-83 (Fig. 3, J and K). Consistent with reduced fatty acid release into the media, VAMP7 and syntaxin 4 knockdown neurons transported less Red-C12 to glia when co-cultured on separate coverslips (Fig. 4, A-D) and accumulated more Red-C12-positive puncta (Fig. 4, E-H).

**Figure 3.**
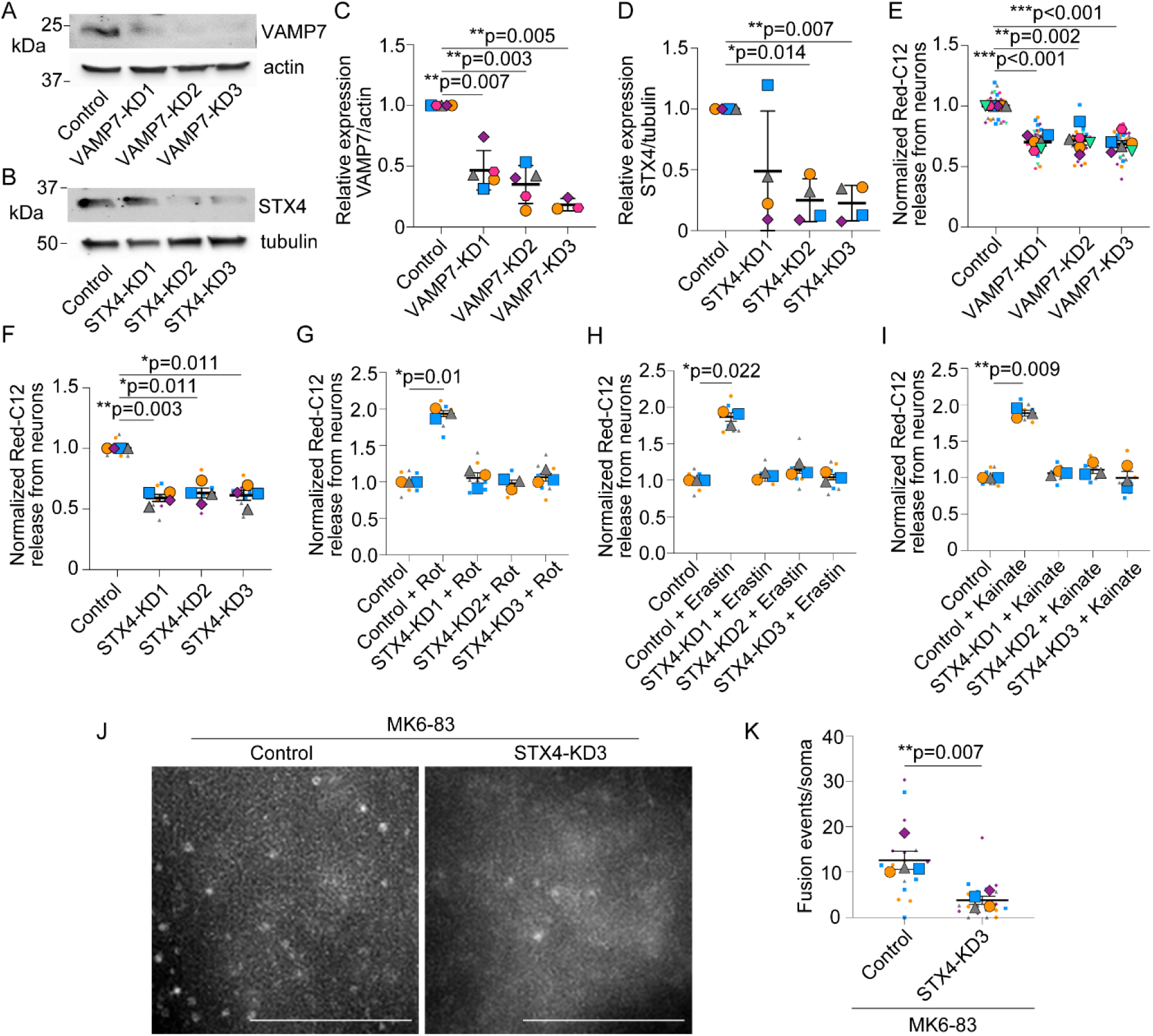
VAMP7 or syntaxin 4 knockdown decreases fatty acid release from neurons. (A) Neurons were transduced with lentivirus expressing non-targeting shRNAmiR (control) or VAMP7 targeting shRNAmiRs (KD1-3). Cell lysates were analyzed by Western blot for VAMP7 levels and β-actin. (B) Neurons were transduced with lentivirus expressing non-targeting shRNAmiR (control) or syntaxin 4 (STX4) targeting shRNAmiRs (KD1-3). Cell lysates were analyzed by Western blot for STX4 levels and β3-tubulin. (C) Levels of VAMP7/β-actin were normalized to non-targeting control. n = 3-5 independent experiments; mean ± SD. One sample t-test with Bonferroni correction. (D) Levels of STX4/β3-tubulin were normalized to non-targeting control. n = 4 independent experiments; mean ± SD. One sample t-test with Bonferroni correction. (E) Neuron-conditioned HBSS analyzed for Red-C12 fluorescence and normalized to non-targeting control neurons. n = 6 independent experiments; mean ± SEM. One sample t-test with Bonferroni correction. (F) Neuron-conditioned HBSS analyzed for Red-C12 fluorescence and normalized to non-targeting control neurons. n = 4 independent experiments; mean ± SEM. One sample t-test with Bonferroni correction. (G-I) Neuron-conditioned media analyzed for Red-C12 fluorescence and normalized to non-targeting shRNAmiR control neurons. n = 3 independent experiments, mean ± SEM. One sample t-test with Bonferroni correction. (J) Maximum intensity projection of syntaxin 4 or non-targeting control neurons treated with MK6-83 and imaged by TRIF microscopy for 30 seconds. Scale bars, 5 ∝m. (K) The number of fusion events imaged by TIRF microscopy and normalized to soma size. n = 4 independent experiments; mean ± SEM. Student’s t-test.

**Figure 4.**
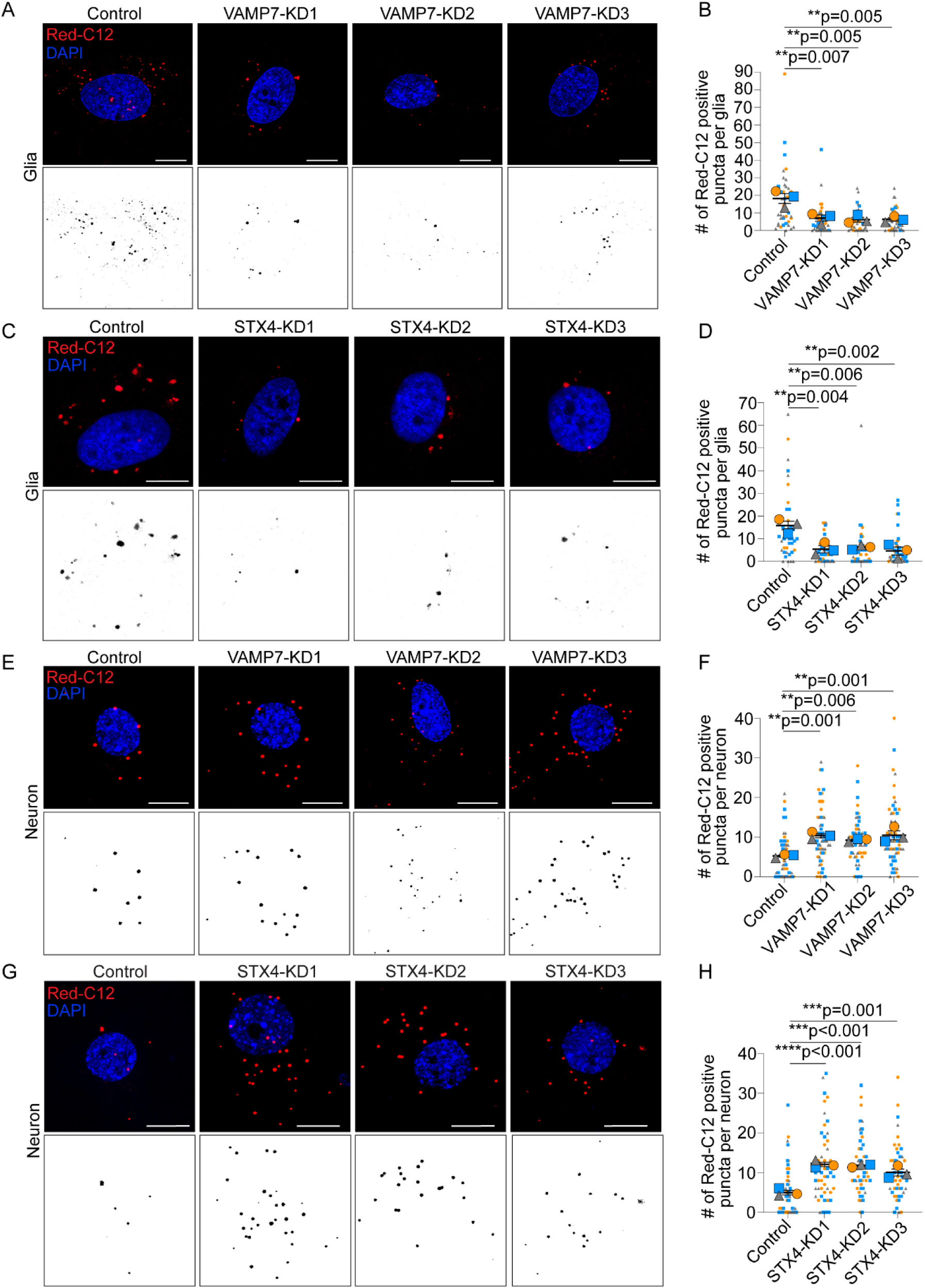
Neuronal VAMP7 or syntaxin 4 knockdown decreases fatty acid transfer to glia. (A and C) Cropped confocal image of glia post-transfer assay in HBSS. Scale bars, 10 ∝m. (B and D) Red-C12 positive lipid droplets in glia following fatty acid transfer assay n = 3 independent experiments; mean ± SEM. One-way ANOVA with Dunnett’s post test. (E and G) Confocal maximum intensity projection of neurons loaded with Red-C12. Scale bars, 10 ∝m. (F and H) The number of Red-C12 positive puncta per neuron. n = 3 independent experiments; mean ± SEM. One-way ANOVA with Dunnett’s post test.

We next turned to a *Drosophila* model system to test whether lysosomal exocytosis contributes to fatty acid transport and glial lipid droplet formation in animals. We induced oxidative stress in photoreceptor neurons by using an *Rh:ND42 RNAi* transgene which expresses RNAi targeting the mitochondrial complex I gene, *ND42,* under the control of a photoreceptor-specific *ninaE* promoter (*Rh*) (Liu et al., 2015). This induces lipid droplet formation in the surrounding pigment glia (Fig. 5, A-C). We then knocked down endogenous drosophila d*VAMP7* or *dSTX4* in these flies using two independent *UAS-RNAi* fly lines for each gene. Knockdown efficiency was confirmed in both larvae and adult flies when *UAS-RNAi* were expressed using the ubiquitous driver *daughterless-GAL4 (da-GAL4)* (Fig. S2, A-D). We then examined whether decreased expression of d*VAMP7* or *dSTX4* affects glial lipid droplet formation in a neuron- or glial-specific manner. Consistent with our *in vitro* results, knockdown of d*VAMP7* or *dSTX4* in photoreceptor neurons using the *rhodopsin-GAL4* driver (*Rh1-GAL4*) significantly decreased glial lipid droplets (Fig. 5, A and D). This effect was cell-type specific as knockdown in surrounding pigment glia using the *54C-GAL4* driver had no effect on glial lipid droplet numbers (Fig. 5, A and E). No glial lipid droplets were observed in the absence of oxidative stress (Fig. S2, E-G). Altogether, these results show that during oxidative stress, lysosomal exocytosis contributes to fatty acid release from neurons.

**Figure 5.**
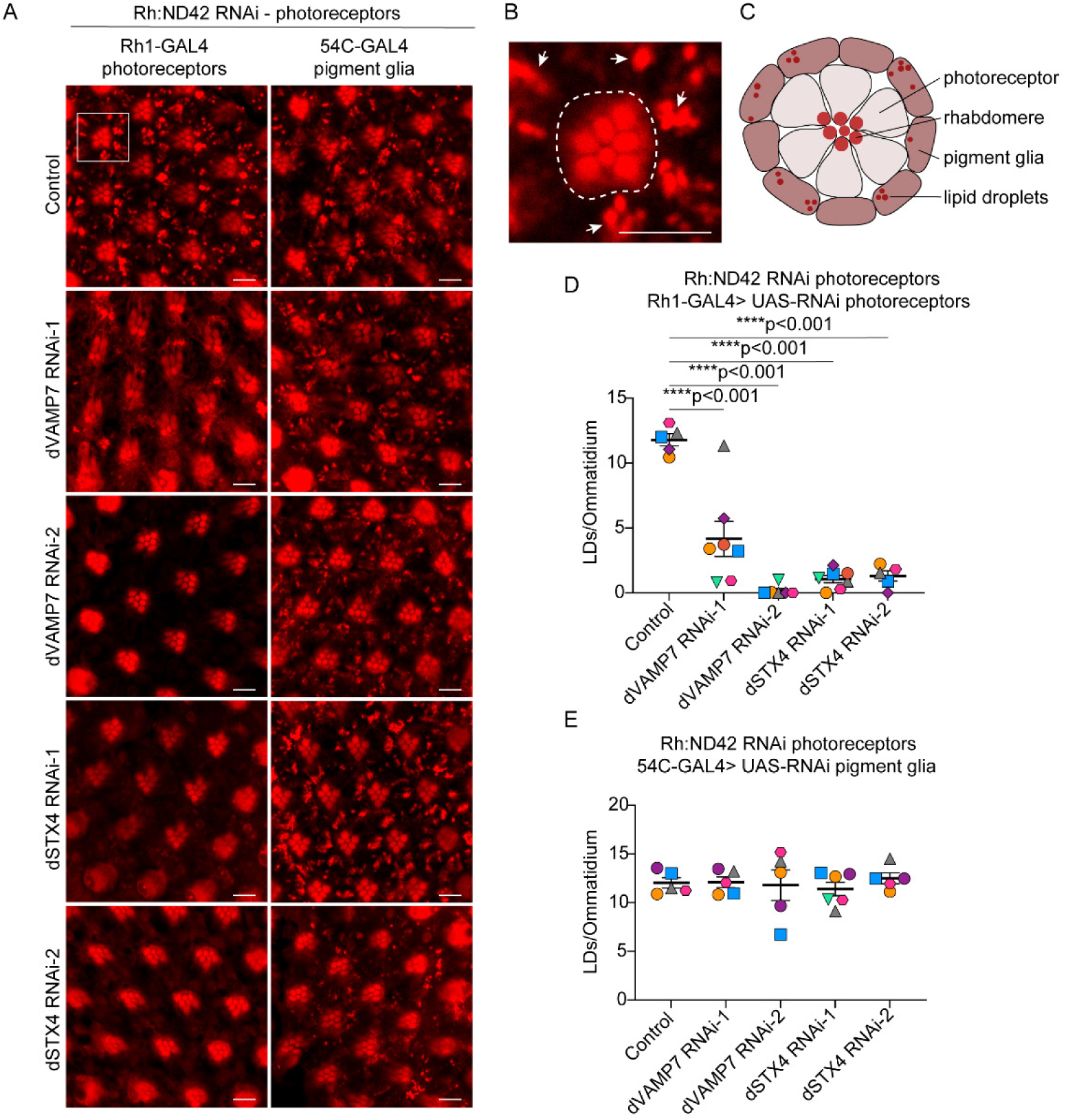
*In vivo* glial lipid droplet formation in the presence of ROS requires neuronal VAMP7 and syntaxin 4. (A and B) Confocal image of Nile Red positive lipid droplets (LD) in pigment glial cells upon ROS induction in photoreceptor neurons via expression of *ND42* RNAi under the control of the *ninaE (Rh)* promoter. *UAS-RNAi* directed against d*VAMP7* and d*STX4* genes was induced in neurons using *Rh1-GAL4* and in pigment glia using *54C-GAL4*. Control fly is UAS-empty. (B) Boxed area is shown as magnified. Photoreceptors (dashed lines) do not accumulate LDs while pigment glia (arrowheads) do. Scale bars, 5 ∝m. (C) Schematic of *Drosophila* ommatidium of the retina. (D) Number of lipid droplets per ommatidium when d*VAMP7* and d*STX4* is knocked down in photoreceptor neurons (*Rh1-GAL4*). Minimum n = 5 animals per genotype; mean ± SEM. One-way ANOVA with Tukey’s post test. (E) Number of lipid droplets per ommatidium when d*VAMP7* and d*STX4* is knocked down in pigment glia (*54C-GAL4*). Minimum n = 5 animals per genotype; mean ± SEM. One-way ANOVA with Tukey’s post test.

### Degradation is not required for autolysosome exocytosis

Cells increase lipid droplet formation during multiple forms of cellular stress including starvation and oxidative stress (Ralhan et al., 2021). One source of fatty acids that fuel lipid droplet growth under these conditions is degradation of membranes and organelles via autophagy (Ioannou, Jackson, et al., 2019; Rambold et al., 2015). We directly tested whether fatty acids generated by autophagy are released from neurons as previously suggested (Ioannou, Jackson, et al., 2019; Rambold et al., 2015). As expected, inhibiting autophagosome formation using 3-methyladenine (3-MA) decreases Red-C12 accumulation in neurons (Fig. 6, A and B) and Red-C12 release into the media (Fig. 6 C). The ability of 3-MA to block autophagy in neurons was evidenced by decreased LC3-II by western blotting in the presence of chloroquine compared to chloroquine alone (Fig. S3, A and B). These data indicate that autophagy supplies lipids released by neurons.

**Figure 6.**
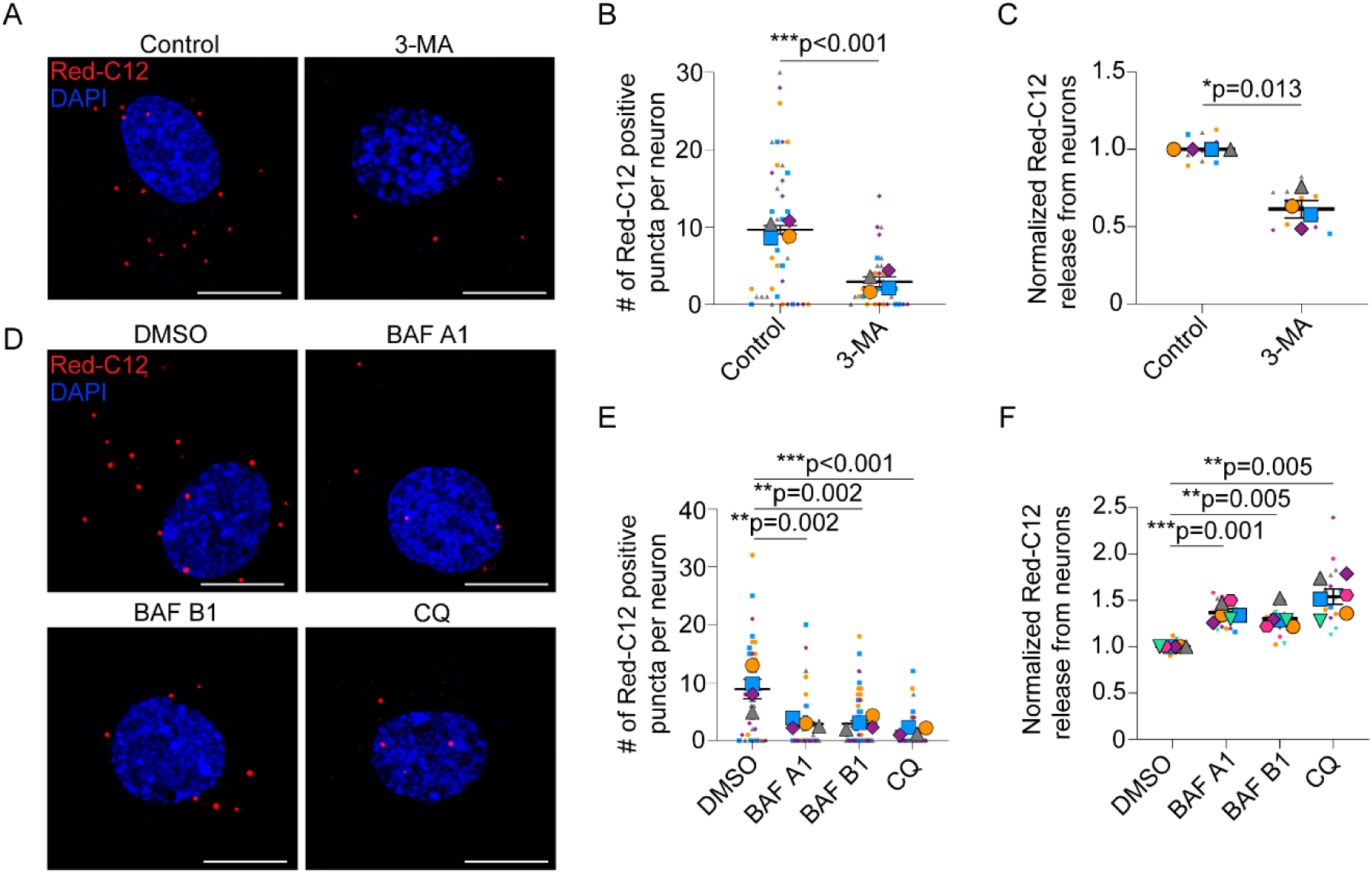
Neurons release autophagy-derived lipids independent of degradation. (A) Confocal maximum intensity projection of neurons loaded with Red-C12 and treated with HBSS ± 3-methyladenine (3-MA). Scale bars, 10 ∝m.(B) Number of Red-C12 positive puncta per neuron. n = 4 independent experiments; mean ± SEM. Student’s T-test. (C) Neuron-conditioned HBSS analyzed for Red-C12 fluorescence normalized to control neurons. n = 4 independent experiments; mean ± SEM. One sample t-test with Bonferroni correction. (D) Confocal maximum intensity projection of neurons loaded with Red-C12 and treated with in HBSS ± DMSO, bafilomycin A1 (BAF A1), bafilomycin B1 (BAF B1), or chloroquine (CQ). Scale bars, 10 ∝m.(E) Number of Red-C12 positive puncta per neuron. n = 4 independent experiments; mean ± SEM. One-way ANOVA with Dunnett’s post test. (F) Neuron-conditioned HBSS analyzed for Red-C12 fluorescence and normalized to DMSO treated neurons. n = 6 independent experiments; mean ± SEM. One sample t-test with Bonferroni correction.

We then tested whether autophagic degradation of membranes and/or organelles is required for release using drugs that target later stages of autophagy. Bafilomycin A1 (Baf A1), B1 (Baf B1) and chloroquine inhibit lysosomal acidification thereby decreasing its degradative function, as well as inhibiting fusion of autophagosomes with lysosomes (Mauthe et al., 2018; Mauvezin & Neufeld, 2015). The effectiveness of these drugs to inhibit late stages of autophagy in neurons was evidenced by the increased LC3-II by western blotting (Fig. S3, A and B). Similar to blocking early autophagy, Baf A1, Baf B1 and chloroquine decreased Red-C12 accumulation in neurons (Fig. 6, D and E). This could be explained by decreased degradation of membranes into free fatty acids. However, unlike blocking early stages of autophagy, Baf A1, Baf B1 and chloroquine stimulated lipid release (Fig. 6 F). This may be attributed to the drugs ability to increase localized calcium levels thereby stimulating exocytosis (Buratta et al., 2020; Camacho et al., 2008). Altogether these data indicate neurons exocytose lysosomes containing autophagy-derived lipids, also known as autolysosomes, however, degradation of contents may not be required.

### Autolysosomal exocytosis rids neurons of peroxidated fatty acids and iron

We next looked at the role of autolysosomal exocytosis in removing peroxidated lipids from neurons. HBSS treatment renders neurons vulnerable to oxidative stress and lipid peroxidation. This is indicated by increased CellRox Green staining (Fig. S1, C and D) and by BODIPY 581/591 (BD-C11) staining, a ratiometric sensor that shifts its fluorescence emission peak from 590 (red) to 510 (green) nm upon lipid peroxidation (Fig. S3, C and D). Under these conditions, neuronal lysosomes labelled with LysoTracker Blue frequently contain peroxidated lipids stained with BD-C11 within neurites (Fig. 7, A and B), but less often in the soma (Fig. S3 E).

**Figure 7.**
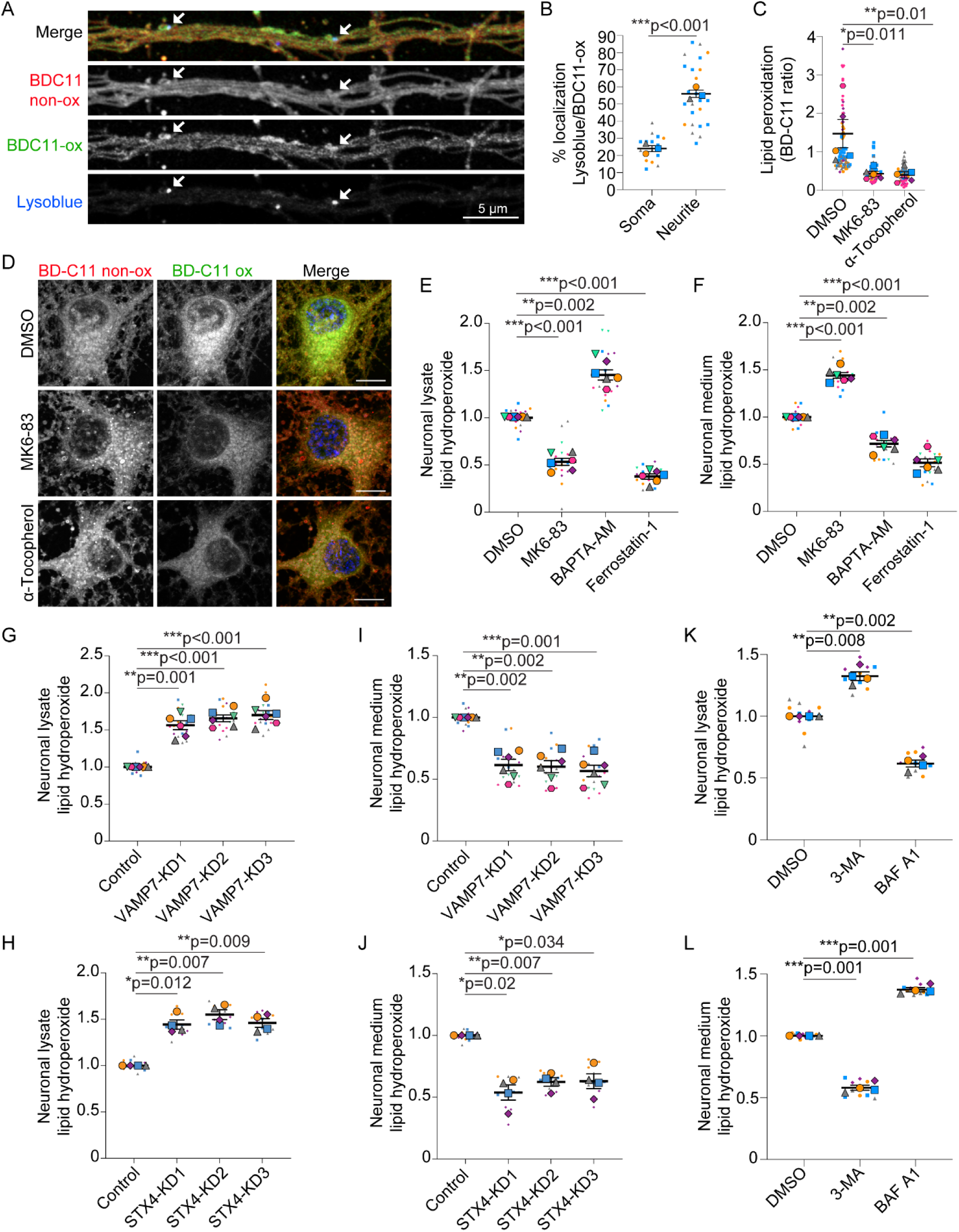
Autolysosomal exocytosis rids neurons of peroxidated fatty acids. (A) Airyscan live-cell image of a neurite in HBSS labelled with LysoTracker Blue and BD-C11. non-peroxidated lipid (BD-C11 non-ox) and peroxidated lipids (BD-C11-ox). Arrows highlight co-localization. Scale bar, 5 ∝m. (B) Co-localization between peroxidated lipids (BD-C11-ox) and LysoTracker Blue in neurons treated in HBSS. n = 3 independent experiments; mean ± SEM. Student’s t-test. (C and D) Confocal image of neurons displayed as maximum intensity projections HBSS ± DMSO, MK6-83 or α-tocopherol labelled with BD-C11. n = 5 independent experiments; mean ± SEM. One-way ANOVA with Dunnett’s post test. Scale bars, 10 ∝m. (E and F) Neuronal lysates (E) and neuron-conditioned HBSS (F) were analyzed for lipid hydroperoxides and normalized to DMSO treated neurons. n = 6 independent experiments; mean ± SEM. One sample t-test with Bonferroni correction. (G and I) Neuronal lysates (G) and neuron-conditioned HBSS (I) were analyzed for lipid hydroperoxides and normalized to non-targeting control neurons. n = 6 independent experiments; mean ± SEM. One sample t-test adjusted with Bonferroni correction. (H and J) Neuronal lysates (H) and neuron-conditioned HBSS (J) were analyzed for lipid hydroperoxides and normalized to non-targeting control neurons. n = 4 independent experiments; mean ± SEM. One sample t-test with Bonferroni correction. (K and L) Neuronal lysates (K) and neuron-conditioned HBSS (L) were analyzed for lipid hydroperoxides and normalized to DMSO treated neurons. n = 4 independent experiments; mean ± SEM. One sample t-test with Bonferroni correction.

We then assessed whether neuronal lipid peroxidation is affected by stimulating exocytosis with MK6-83. We observed reduced BD-C11 ratio (green to red) indicative of decreased lipid peroxidation when exocytosis was stimulated with MK6-83 (Fig. 7, C and D). The reduction in lipid peroxidation was comparable to treatment with α-tocopherol, a lipophilic antioxidant (Fig. 7, C and D). Next, endogenous lipid hydroperoxides were extracted from neuronal lysates and neuron-conditioned media and reacted with ferrous ions to produce ferric ions that were detected using thiocyanate ions as a chromogen. As a control, we treated neurons with ferrostatin-1, a lipophilic antioxidant and potent inhibitor of ferroptosis, and saw reduced lipid hydroperoxides in both lysates and conditioned-medium (Fig. 7, E and F). Stimulating exocytosis with MK6-83 decreased lipid hydroperoxides in neurons (Fig. 7 E) and increased lipid hydroperoxide release into media (Fig. 7, F). Conversely, blocking autolysosomal exocytosis with VAMP7 knockdown, syntaxin 4 knockdown or BAPTA-AM treatment increased lipid hydroperoxides in neuronal lysates and decreased lipid hydroperoxide released into the media (Fig. 7, E-J).

Since autolysosomes contain peroxidated lipids, we next tested if inhibiting autophagy prevents release of lipid hydroperoxides. Indeed, early autophagy inhibitor 3-MA caused lipid hydroperoxide accumulation in neurons with a concomitant decrease in lipid hydroperoxides released (Fig. 7, K and L). BAF A1 on the other hand, which prevents degradation and stimulates exocytosis, relieves the lipid hydroperoxide burden in neurons and increases its release into the media (Fig. 7, K and L). Collectively, these results indicate that neurons release peroxidated fatty acids by autolysosomal exocytosis and that blocking this release pathway causes an accumulation of lipid hydroperoxides.

Since lipid peroxidation is an iron-dependent process, and lysosomes are central to iron metabolism (U. T. Brunk, 1989; Gray & Woulfe, 2005), we tested whether autolysosomal exocytosis also affects total iron levels (Fe^2+^ and Fe^3+^). Similar to lipid hydroperoxides, inhibiting autophagy with 3-MA or autolysosomal exocytosis with VAMP7 or syntaxin 4 knockdown increased iron accumulation in neuronal lysates (Fig. 8, A-C) and decreased iron released into the media (Fig. 8, D-F). This indicates that like lipid hydroperoxides, iron is also released by neurons via autolysosomal exocytosis and that blocking this pathway causes an accumulation of iron.

**Figure 8.**
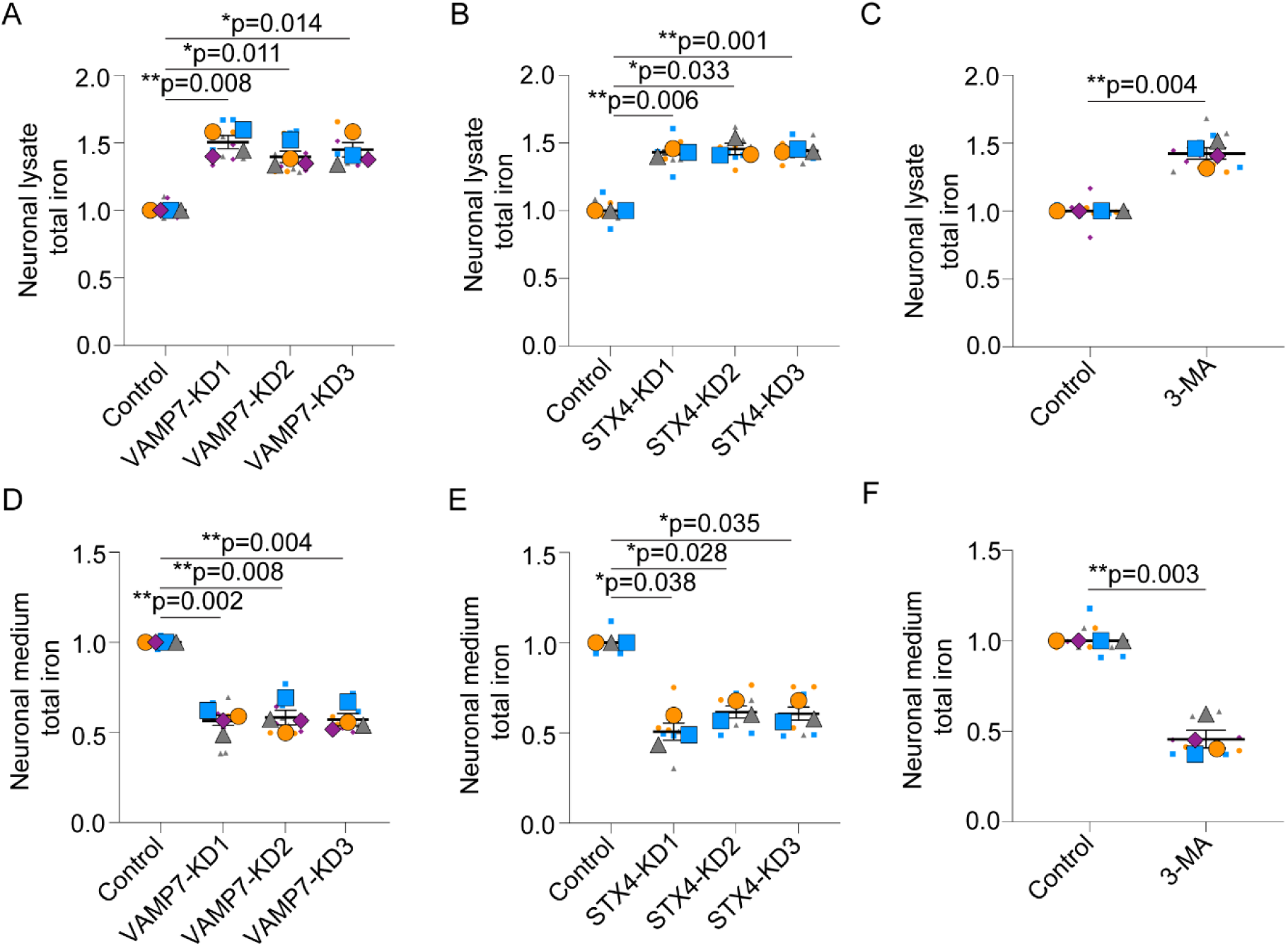
Autolysosomal exocytosis promotes iron release from neurons. (A and D) Neuronal lysates (A) and neuron-conditioned HBSS (D) were analyzed for total iron. VAMP7 knockdown neurons were normalized to non-targeting control neurons. n = 4 independent experiments; mean ± SEM. One sample t-test with Bonferroni correction. (B and E) Neuronal lysates (B) and neuron-conditioned HBSS (E) were analyzed for total iron. STX4 knockdown neurons were normalized to non-targeting control neurons. n = 3 independent experiments; mean ± SEM. One sample t-test with Bonferroni correction. (C and F) Neuronal lysates (C) and neuron-conditioned HBSS (F) were analyzed for total iron. 3-methyladenine (3-MA) treated neurons were normalized to control neurons. n = 4 independent experiments; mean ± SEM. One sample t-test with Bonferroni correction.

### Autolysosomal exocytosis protects neurons from ferroptosis

Accumulation of iron and/or lipid hydroperoxides have been implicated in lysosomal dysfunction (Jahng et al., 2019; Krohne et al., 2010). Consistently, syntaxin 4 knockdown decreased activity of cathepsin D, a lysosomal protease commonly used to assess lysosomal function (Fig. 9 A). Therefore, blocking autolysosomal exocytosis results in dysfunctional lysosomes, presumably due in part to the accumulation of iron and/or lipid hydroperoxides.

**Figure 9.**
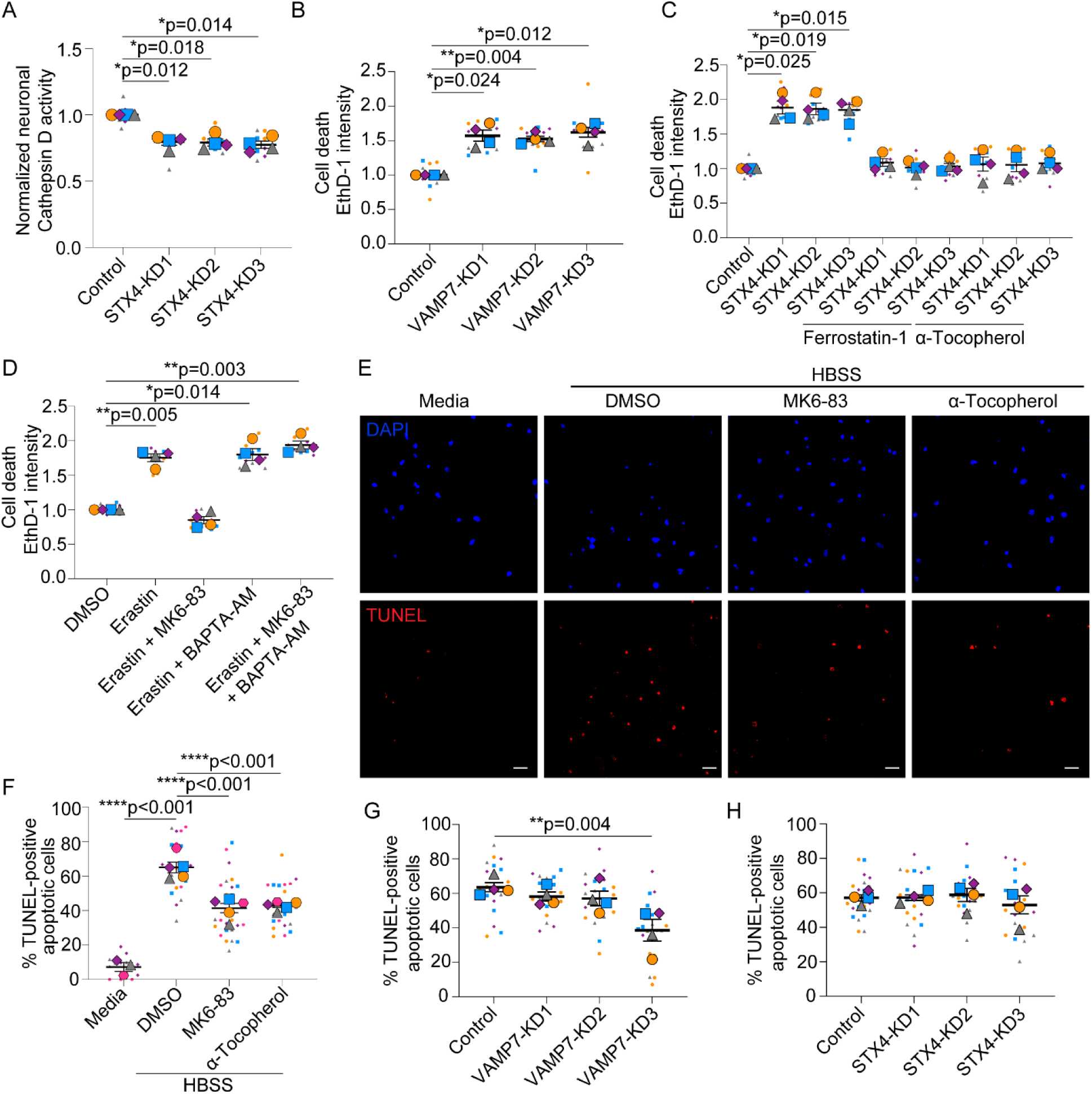
Autolysosomal exocytosis protects neurons from ferroptosis. (A) Neurons were transduced with lentivirus expressing non-targeting shRNAmiR (control) or shRNAmiRs targeting syntaxin 4 (STX4), analyzed for cathepsin D activity and normalized to non-targeting control neurons. n = 4 independent experiments; mean ± SEM. One sample t-test with Bonferroni correction. (B and C) Neurons were transduced with lentivirus expressing non-targeting shRNAmiR (control) or shRNAmiRs targeting VAMP7 or syntaxin 4 (STX4), treated with HBSS ± DMSO, ferrostatin-1 or α-tocopherol, stained with ethidium homodimer-1 (EthD-1), and normalized to non-targeting control neurons. n = 4 independent experiments; mean ± SEM. One sample t-test with Bonferroni correction. (D) Neurons were treated with media ± DMSO, erastin, MK6-83 or BAPTA-AM and stained with EthD-1 and normalized to DMSO treated neurons. n = 4 independent experiments; mean ± SEM. One sample t-test with Bonferroni correction. (E and F) Confocal images neurons in media or HBSS ± DMSO, MK6-83 or α-tocopherol. Apoptotic cells were labelled with TUNEL. Scale bars, 25 ∝m. n = 3-5 independent experiments. Mean ± SEM. One-way ANOVA with Tukey’s post test. (G and H) Neurons were transduced with lentivirus expressing shRNAmiR as above, treated with HBSS and apoptotic cells were labelled with TUNEL. n = 4 independent experiments; mean ± SEM. One-way ANOVA with Dunnett’s post test.

Accumulation of peroxidated lipids and iron also triggers ferroptosis, a biochemically distinct form of cell death (Dixon et al., 2012; Li et al., 2020; Ye et al., 2020). We next examined whether autolysosomal exocytosis effects neuronal death by ferroptosis. We began by staining with ethidium homodimer-1 that labels dying cells with disrupted membrane integrity irrespective of the type of cell death. Stimulating exocytosis with MK6-83 protects neurons from non-specific cell death as measured by reduced ethidium homodimer-1 fluorescence (Fig. S4, A and B). Reductions in cell death were comparable to treatment with lipophilic antioxidants α-tocopherol and ferrostatin-1 (Fig. S4 A). Conversely, VAMP7 or syntaxin 4 knockdown increased ethidium homodimer-1 fluorescence in HBSS (Fig. 9, B and C), revealing a protective function for the autolysosomal exocytosis pathway.

To tease apart the contribution of oxidative stress from starvation, we also measured cell death of neurons in complete media when ferroptosis was selectively induced using erastin (Dixon et al., 2014). Here, cell death induced by erastin was rescued by MK6-83 to induce exocytosis (Fig. 9, D) and exacerbated by syntaxin 4 knockdown (Fig. S4, C). BAPTA-AM, a calcium chelator, prevents MK6-83 from reducing erastin-mediated cell death, (Fig. 9, D and S4, B). This calcium dependence further strengthens the conclusion that the protective effects of MK6-83 are mediated by exocytosis. Cell death induced by syntaxin 4 knockdown, in either HBSS or complete media, can be rescued with lipophilic antioxidants α-tocopherol and ferrostatin-1 (Fig. 9, C and S4, C), further indicating the importance of peroxidated lipids in neuronal death caused by blocking exocytosis.

Since there is considerable crosstalk between ferroptosis and apoptosis, we next looked at whether exocytosis affects apoptosis. Terminal deoxynucleotidyl transferase dUTP nick end labelling (TUNEL) stains fragmented DNA specific to apoptosis (Kyrylkova et al., 2012), and is not observed in ferroptosis (Dixon et al., 2012). HBSS treatment increased apoptotic cell death as measured by TUNEL staining (Fig. 9, E and F). Stimulating exocytosis with MK6-83 reduced apoptotic cell death to levels comparable to α-tocopherol treatment (Fig. 9, E and F). However, VAMP7 or syntaxin 4 knockdown had no affect on TUNEL staining (Fig. 9, G and H). Thus, autolysosomal exocytosis appears to have only a modest role in protecting cells from apoptosis. Instead, autolysosomal exocytosis protects neurons largely from ferroptosis.

### Neurons release lipid hydroperoxides in lipid-protein particles

Our results showing the protective function of neuronal fatty acid release led us to ask what species of lipids are released. In addition to Red-C12, neuronal release of BODIPY cholesteryl ester (BD-CE) is increased by stimulating exocytosis with MK6-83 and decreased by inhibiting exocytosis with BAPTA-AM or VAMP7 knockdown (Fig. 10, A and B). However, Red-C12 could remain soluble as a fatty acid bound to carrier proteins, and both Red-C12 and BD-CE could exist as phospholipids or neutral lipids and incorporated into membranes or lipoprotein-particles (Quinlivan et al., 2017). MK6-83 and BAPTA-AM had no effect on the release of the neutral lipid stain BD493 (Fig. S5 A), suggesting it is unlikely that neutral lipids are released from neurons through an exocytosis pathway.

**Figure 10.**
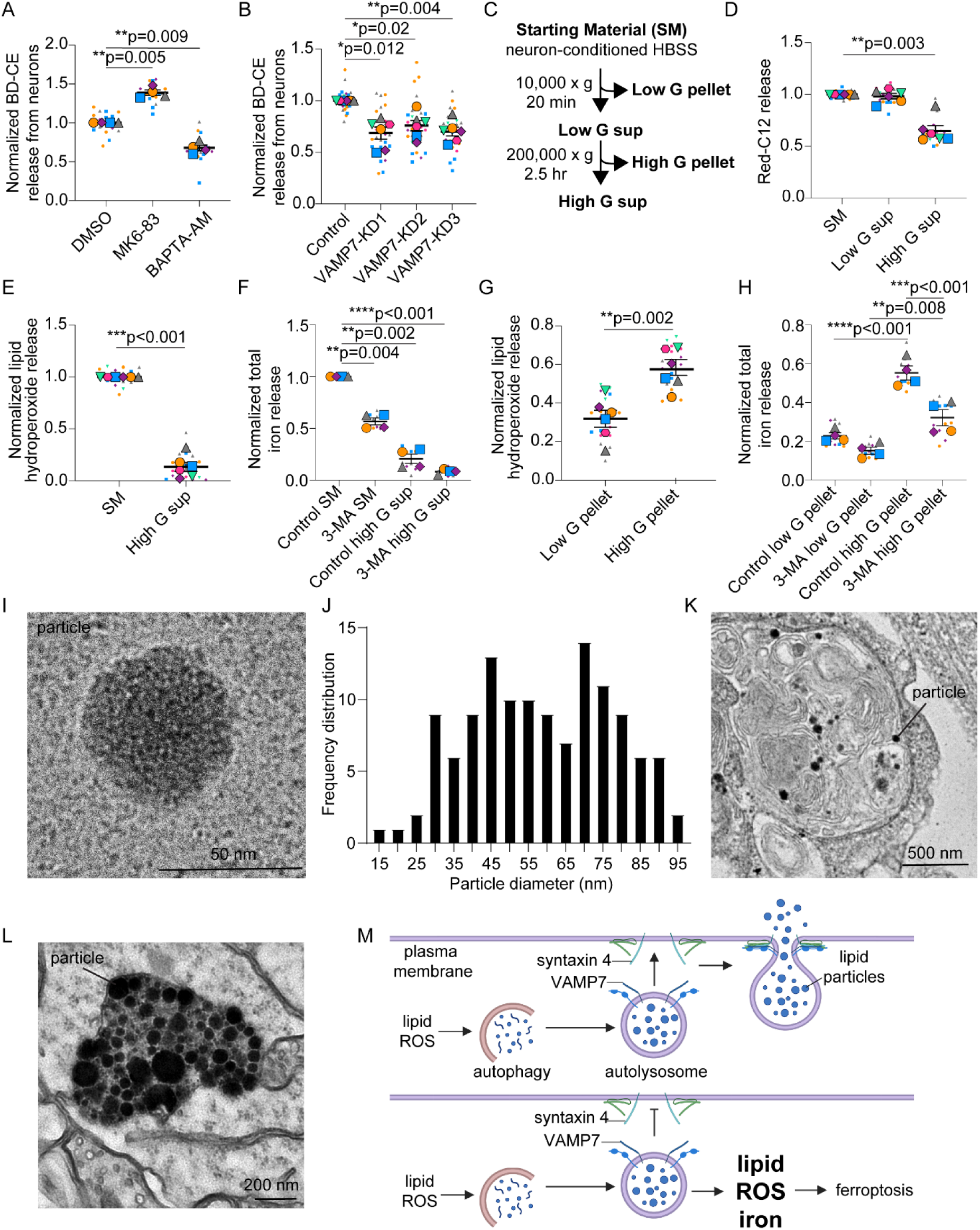
Neurons release peroxidated lipids and iron as lipid-protein particles. (A) Neuron-conditioned HBSS analyzed for BODIPY-cholesteryl ester (BD-CE) fluorescence and normalized to DMSO treated neurons. n = 4 independent experiments; mean ± SEM. One sample t-test adjusted with Bonferroni correction. (B) Neurons were transduced with lentivirus expressing non-targeting shRNAmiR (control) or shRNAmiRs targeting VAMP7. Neuron-conditioned HBSS analyzed for BD-CE fluorescence. n = 6 independent experiments; mean ± SEM. One sample t-test with Bonferroni correction. (C) Schematic of centrifugation assay. Supernatant (sup). (D) Neuron-conditioned HBSS analyzed for Red-C12 fluorescence following depletion by centrifugation. n = 6 independent experiments; mean ± SEM. One sample t-test with Bonferroni correction. (E) Neuron-conditioned HBSS analyzed for lipid hydroperoxides following depletion by centrifugation. n = 6 independent experiments; mean ± SEM. One sample t-test with Bonferroni correction. (F) Neurons were treated with HBSS ± 3-methyladenine (3-MA) and neuron-conditioned HBSS was analyzed for total iron following depletion by centrifugation. n = 4 independent experiments; mean ± SEM. One sample t-with Bonferroni correction. (G) Pellets obtained from centrifugation of neuron-conditioned HBSS were analyzed for lipid hydroperoxides. n = 6 independent experiments; mean ± SEM. Student’s T-test. (H) Pellets obtained from centrifugation of neuron-conditioned HBSS were analyzed for total iron. n = 4 independent experiments; mean ± SEM. One-way ANOVA with Dunnett’s post test. (I) Lipid-protein particle from high G centrifugation of neuron-conditioned HBSS imaged by cryoEM. Scale bar, 50 nm. (J) Frequency distribution of lipid-protein particle diameter determined by cryoEM. (K) Cultured neurons stained with imidazole-buffered osmium were imaged by transmission electron microscopy. Scale bar, 500 nm. (L) Hippocampal neurons from sectioned mouse brain, stained with imidazole-buffered osmium were imaged by transmission electron microscopy. Scale bar, 200 nm. (M) Schematic of autolysosomal exocytosis of lipids from neurons.

To further explore what neurons are releasing by autolysosomal exocytosis, we used differential centrifugation to deplete components from neuron-conditioned HBSS and evaluated the levels of Red-C12, lipid hydroperoxides, or iron remaining (Fig. 10 C). Low-G centrifugation to deplete apoptotic bodies had no affect on the levels of Red-C12 remaining in the supernatant, whereas high-G centrifugation reduced Red-C12 by nearly half (Fig. 10 D). This indicates that fatty acids are released as components of the high-G pellet in addition to a soluble and/or buoyant pool of fatty acids (Ioannou, Jackson, et al., 2019). Lipid hydroperoxides and iron, however, are largely depleted from neuron-conditioned medium following high-G centrifugation (Fig. 10, E and F and Fig. S5 B). Both lipid hydroperoxides and iron are concentrated in the high-G pellet (Fig. 10, G and H and Fig. S5 C).

To determine what lipid-containing structures are present in the high-G pellet, we performed cryoEM to visualize components of the high-G pellet in their close-to-native hydrated states. We discovered electron-dense particles (Fig. 10, I and S5 D), ranging in diameter from 15-95 nm (Fig. 10 J). Given their presence in the high-G supernatant indicating they are heavy, and electron-dense appearance by cryoEM, we speculate these are lipid-protein particles found within residual bodies. Residual bodies, autolysosomes containing partially digested or undigested lipids, are precursors to lipofuscin which is common in aged neurons. They are enriched in peroxidated lipids and iron. Consistently, we observed similar lipid-protein particles by transmission electron microscope in autophagosomes and/or autolysosomes in cultured neurons (Fig. 10 K and S5 E), while in the adult mouse brain, they are in electron-dense vesicles resembling lipofuscin (Fig. 10 L and S5 F).

The high-G pellet also contains extracellular vesicles as expected (Fig. S5 D) (Matthies et al., 2020). Fusion of multivesicular bodies to release exosomes could utilize the same SNARE machinery, VAMP7 and syntaxin 4, for their release. However, given that 3-MA stimulates exosome release (Fig. S5, G and H) (Wu et al., 2021), yet decreases lipid hydroperoxide and iron release (Fig. 7, L and 8, F), we reason that exosomes may play only a minor role in protecting neurons from lipid peroxidation. Like lipid hydroperoxides, iron release and enrichment in the high-G pellet is decreased with 3-MA treatment (Fig. 10, F and H). Our data reveal autolysosomal exocytosis of lipid-protein particles as a likely mechanism for riding neurons of excess peroxidated lipids and iron.

## DISCUSSION

Lipid release from neurons has emerged as an important feature during oxidative stress. Here, we identified autolysosomal exocytosis as a key pathway for neuronal lipid release. We show that autolysosomal exocytosis removes peroxidated lipids and iron from neurons and protects them from cell death by ferroptosis. We also identified the lipid-containing structures that likely facilitate this process; that being lipid-protein particles. As discussed below, autolysosomal exocytosis protects neurons from lipid peroxidation during oxidative stress and has far reaching implications for numerous neurodegenerative diseases and brain injury.

Lysosomal exocytosis is mediated by the interaction between the lysosomal v-SNARE VAMP7/TI-VAMP and the plasma membrane t-SNAREs syntaxin 4 and SNAP-23 (Rao et al., 2004). We discovered that VAMP7 and syntaxin 4 are needed for lipid release and protect neurons from the damaging effects of lipid peroxidation. VAMP7 also participates in endosomal trafficking events upstream of vesicle fusion with the plasma membrane. This includes fusion of lysosomes with late endosomes (Advani et al., 1999), autophagosomes (Wojnacki et al., 2020), and mitochondrial-derived vesicles (McLelland et al., 2016). Syntaxin 4, being localized to the plasma membrane, facilitates exocytosis with no apparent effect on endosomal transport or autophagy (Takáts et al., 2013). Since targeting either VAMP7 or syntaxin 4 impairs lipid release, this suggests the effects on lipid release are mediated via their exocytic function.

VAMP7 and syntaxin 4 can also facilitate exocytosis of related compartments including autophagosomes, late endosomes and multivesicular bodies (Kimura et al., 2017; Verweij et al., 2018; Wojnacki et al., 2020). Our experiments showing that autophagy supplies peroxidated lipids for release points to exocytosis of the autolysosomes as opposed to late-endosomes or multivesicular bodies. Also, inhibitors of early-stage autophagy stimulate extracellular vesicle release (Wu et al., 2021) but decrease lipid hydroperoxide and iron release further strengthens this conclusion. While the lipid-protein particles in the media resemble the indigestible material that remains in autolysosomes, similar to residual bodies and lipofuscin (U. Brunk & Brun, 1972; Moreno-García et al., 2018), we cannot rule out the contribution of autophagosome secretion in the process (Kimura et al., 2017).

We discovered a critical function for exocytotic release of lipid-protein particles; protection from ferroptosis. During oxidative stress, lipids become peroxidated in the presence of iron and excess ROS. Polyunsaturated phospholipids are the preferred substrate for peroxidation with phosphatidylethanolamines being a prominent phospholipid species that drives ferroptosis (Kagan et al., 2017; Magtanong et al., 2019). Additional lipid species can also undergo oxidative modifications to generate toxic species, such as cholesterol hydroperoxides (Korytowski et al., 2013). Increased lipid peroxidation triggers autophagy to remove damaged membranes that are typically degraded. Our current work indicates autolysosome fusion with the plasma membrane to release their contents prior to complete digestion, to be detoxified by glia as previously shown (Ioannou, Jackson, et al., 2019; Liu et al., 2017). By direct release from these compartments, peroxidated lipids and iron would neither accumulate in the lysosome or re-enter the cytoplasm and cause further damage.

Our study is consistent with genome-wide screens that identified lysosomal defects as a trigger for ferroptosis, specifically in neurons (Tian et al., 2021). Lysosomal dysfunction results in lipofuscin formation, the accumulation of indigested material including peroxidated lipids and iron. This is commonly observed in neurons during aging and in neurodegenerative disease (Moreno-García et al., 2018; Zhou et al., 2017). Exocytosis of autolysosomes (residual bodies) prevents lipofuscin formation in macrophages (Hendriks & Eestermans, 1986). However, this process is not well defined in long-lived cell types such as neurons. Lipofuscin formation in neurons has been attributed to decreased degradative capacity of lysosomes. Our data shows that autolysosomal exocytosis also prevents accumulation of peroxidated lipids, a key component of lipofuscin. We propose that defects in autolysosomal exocytosis may also contribu to lipofuscin formation in neurons in aging and disease.

In cultured neurons, lipids that fail to be released are stored in lipid droplets. This is different from what is observed in aged neurons that rarely form lipid droplets. Neurons cultured from postnatal pups have only recently differentiated from neural progenitor cells and may continue to express genes of neural progenitor cells including those required for lipid droplet formation. Lipid droplets are present in neural stem cells and regulate their ability to proliferate and/or differentiate (Ramosaj et al., 2021). Expression of these genes likely tapers off in fully mature neurons. With no where to store lipids in mature neurons, failure to exocytose lysosomal lipids could then accumulate as lipofuscin, which is rarely observed in young neurons. It would be interesting to explore whether inducing lipid droplet formation in aging neurons could protect lysosomes from dysfunction associated with lipid accumulation.

In addition to exocytic release of lipids, neurons can shed fatty acids using ABCA1 and ABCA7 transporters (Moulton et al., 2021). This raises an important question: How do neurons decide which mechanism of release to use to rid themselves of excess lipids? Both ABCA1 and ABCA7 are largely known for their ability to transport plasma membrane phospholipids and cholesterol to circulating lipoprotein particles (Abe-Dohmae et al., 2006; Hayashi et al., 2005; Wahrle et al., 2004). One appealing possibility is that neurons use ABC transporters on the plasma membrane to constitutively remove lipids during homeostasis while autolysosome exocytosis allows for a bulk release of lipids during oxidative stress. ABCA1 also localizes to late endosomes and ABCA1 defects result in the accumulation of phospholipids and cholesterol in late endosomal compartments (Neufeld et al., 2004). ABCA transporters could similarly affect lipid accumulation in autolysosomes as well. In which case, defects in ABC transports may converge directly on lysosomes with exocytosis being the final step. Another remaining question is which of these pathways does apolipoprotein E (ApoE) act on? ApoE is secreted and acts in concert with ABC transporters to facilitate lipid efflux. Knockdown of ApoE in either a neuron- or glia-specific manner reduces fatty acid release (Ioannou, Jackson, et al., 2019; Liu et al., 2017). This does not preclude the possibility that ApoE might also affect lysosomal exocytosis as the ApoE4 polymorphism associated with Alzheimer’s disease disturbs lysosomal physiology (Fote et al., 2022). Understanding how ApoE fits into these pathways will provide important insight into understanding Alzheimer’s disease.

Finally, the role of extracellular vesicles in mediating lipid release needs to be explored further. While our results indicate that exosomes play only a minor role in removing peroxidated lipids and/or iron from neurons, extracellular vesicles carrying membrane phospholipids, sphingolipids and cholesterol would also contribute to lipid transport between cell types in the brain. We favor a model where different lipid species exit neurons via different pathways and speculate that each has important implications for regulating neuronal health.

## ACKNOWLEDGEMENTS

We thank Peter McPherson for helpful comments on the manuscript. Experiments were performed at the University of Alberta Faculty of Medicine & Dentistry Cell Imaging Core, RRID:SCR_019200. We thank Pinzhang Gao for assistance with EM sample preparation, Monique Copeland for assistance with mouse perfusions and Sarah Hughes, Todd Alexander, Nicolas Touret and Elaine Leslie for sharing critical equipment. We thank Rick Kuo-Jui Huang and Allison Zeher for their support at the NIH IRP CryoEM Consortium. This work utilized the computational resources of the NIH HPC Biowulf cluster (http://hpc.nih.gov).

I.R. is supported by the Canadian Institutes of Health Research (CIHR) Doctoral Award #181551, the Alberta Synergies in Alzheimer’s and Related Disorders program funded by the Alzheimer Society of Alberta and Northwest Territories, the University Hospital Foundation and the Neuroscience and Mental Health Institute, an Alberta Graduate Excellence Scholarship, and a 75^th^ Anniversary Graduate Student Award from the Faculty of Medicine & Dentistry. M.J.M. is supported by the Brain Disorders & Development Fellowship Training Grant from the National Institutes of Health (NIH) T32 NS043124-19. L.D.G. is supported by the Postdoctoral Fellowship Program in Alzheimer’s Disease Research from the BrightFocus Foundation. H.A.P. is supported by the Howard Hughes Medical Institute. H.J.B. is supported by The National Institute on Aging of the NIH R01 AG07326 and U01 AG072439, The Office of Strategic Coordination/Office of the NIH Director U54 NS093793 and R01 HG011795, The Office of Research Infrastructure Programs of the NIH R24 OD022005 and R24 OD031447 and the Jan and Dan Duncan Neurological Research Institute at Texas Children’s Hospital. M.S.I. is supported by a Sloan Research Fellowship from the Alfred P. Sloan Foundation, Canada Research Chairs program, the Heart and Stroke Foundation of Canada (#170722), the Canadian Foundation for Innovation (#40616), and CIHR (#173321).

## AUTHOR CONTRIBUTIONS

I. Ralhan and M.S. Ioannou conceived and designed the study. M.J. Moulton and L.D. Goodman performed and analyzed drosophila experiments. I. Ralhan and G. Plummer performed TIRF imaging. H.J. Bellen supervised drosophila experiments. I. Ralhan and H. Amalia Pasolli performed transmission electron microscopy experiments. D. Matthies performed cryo-electron microscopy experiments. N.Y.J. Lee analyzed the cryo-electron microscopy data. All other experiments and analyses were performed and analyzed by I. Ralhan, J. Chang, and M.S. Ioannou. I. Ralhan and M.S. Ioannou wrote the manuscript with input from all co-authors. M.S. Ioannou supervised the project.

## MATERIALS AND METHODS

**Table 1:**
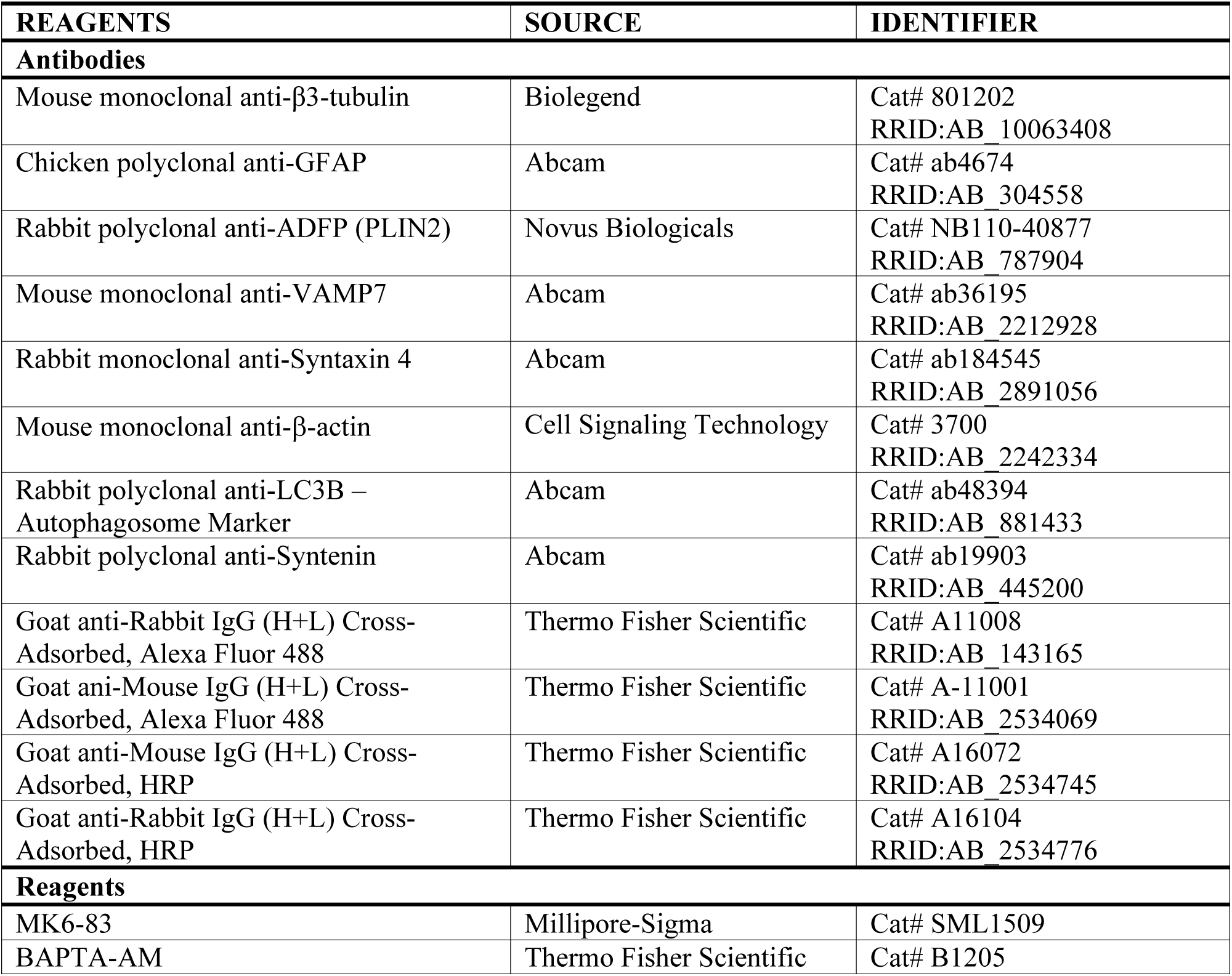
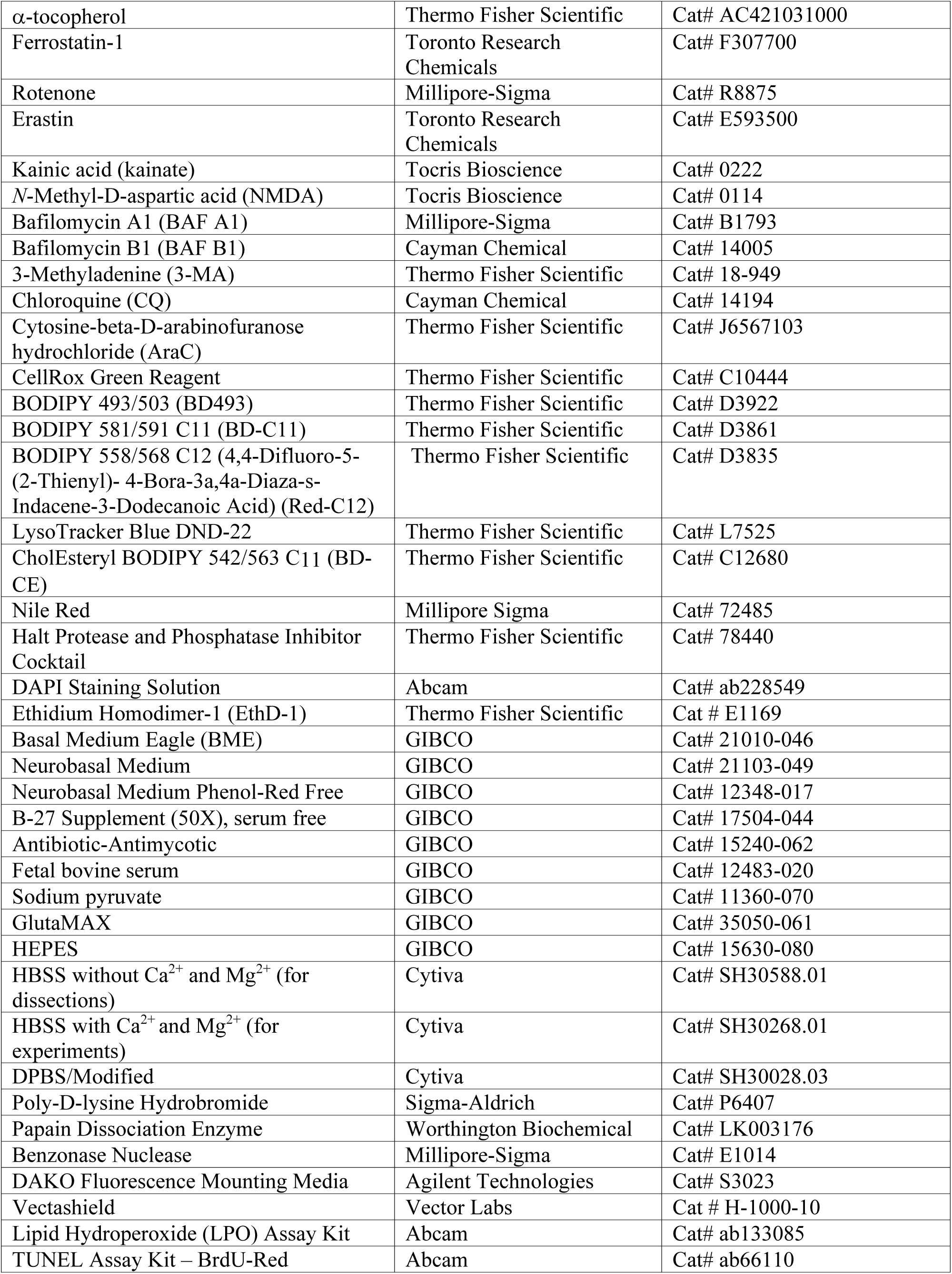
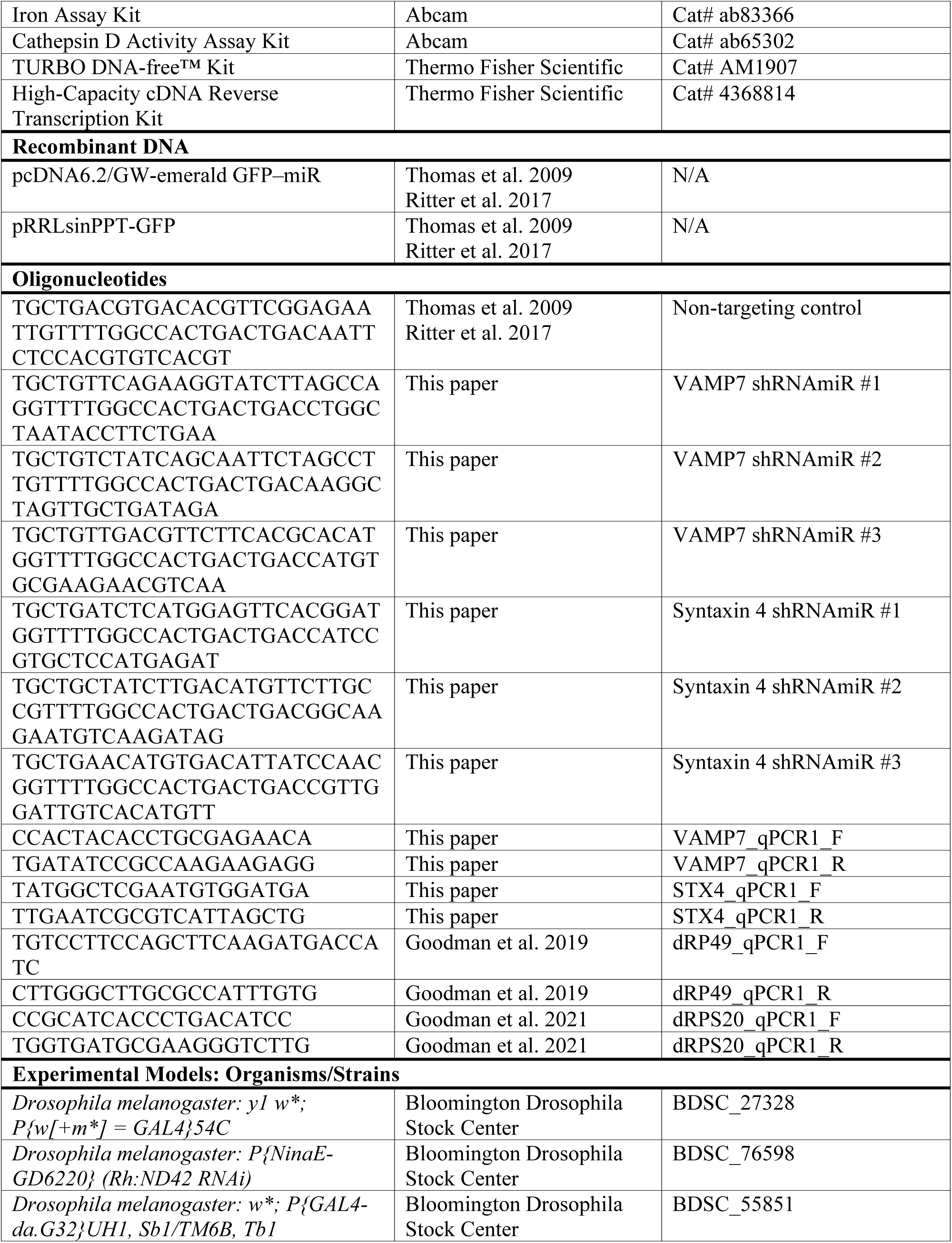
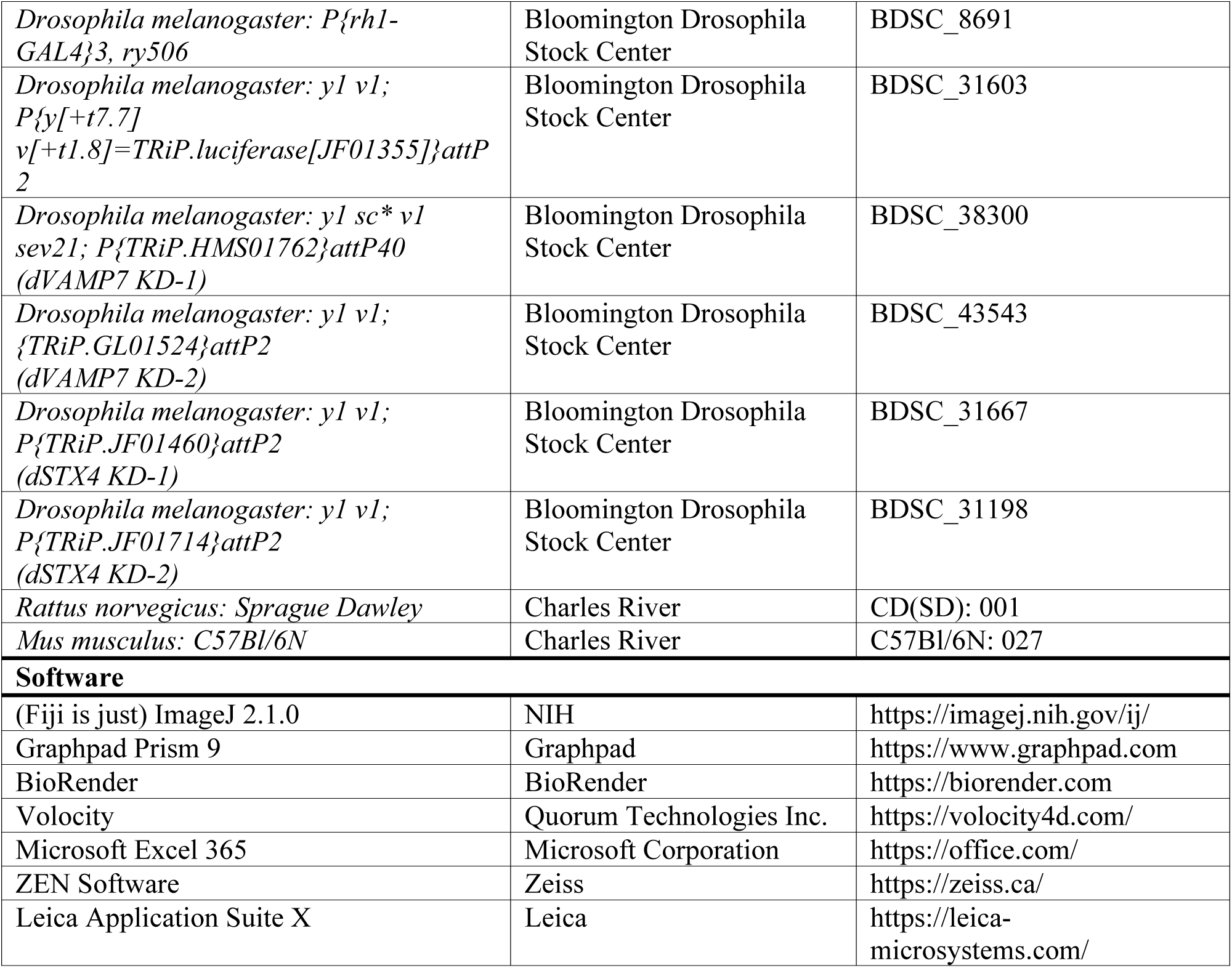
Reagents and Resources.

### Animals

Experiments involving rat cells was approved by and performed in accordance with the Canadian Council of Animal Care at the University of Alberta (AUP#3358). Sprague-Dawley timed pregnant rats were obtained from Charles River Laboratories and arrived at our facility one week prior to birth. Mouse experiments were approved by the Janelia Institutional Animal Care and Use Committee (IACUC 16–146). *Drosophila melanogaster* were raised on standard molasses-based lab diet at 22°C under constant light conditions.

### Primary culture of hippocampal cells

Primary hippocampal cultures were prepared as previously described (Ioannou, Liu, et al., 2019). Briefly, hippocampi were dissected from P0-P1 Sprague-Dawley rat pups and digested with papain and benzonase nuclease. After digestion, the tissues were gently triturated and filtered with a 70-µm nylon cell strainer. Neurons and glia were grown on poly-D-lysine (PDL) coated glass coverslips or plastic tissue culture dishes. Neurons were grown in Neurobasal medium containing B-27 supplement, 2 mM GlutaMax and antibiotic-antimycotic. 1 μM AraC was added to the media 2 days after plating and maintained for 4 days to prevent growth of glia in neuronal cultures. Glia were grown in Basal Eagle Media containing 10% fetal bovine serum, 0.45% glucose, 1 mM sodium pyruvate, 2 mM GlutaMax and antibiotic-antimycotic. All cells were grown at 37°C in 5% CO_2_ and used at DIV 6-9.

### Lentivirus production

pcDNA6.2/GW-emerald GFP–miR, pRRLsinPPT and the pRRLsinPPT-non-targeting-miR control were generous gifts from Dr. Peter McPherson. Target sequences for rat VAMP7 and syntaxin 4 were designed using the BLOCK-iT RNAi Designer (Invitrogen). Target sequences were subcloned first into the pcDNA6.2/GW-emerald GFP–miR cassette and then into the pRRLsinPPT viral expression vector (Ritter et al., 2017). VAMP7-targeting lentiviral particles were generated by VectorBuilder. Syntaxin 4 targeting viral particles were produced in HEK-293T cells as previously described (Ritter et al., 2017; Thomas et al., 2009). In brief, media containing viral particles were collected and filtered with a 0.45-μm filter to remove cell debris and concentrated by centrifugation at 12,000 x g for 15 h at 4°C.

### Lentivirus transduction

Neurons were transduced with lentivirus at DIV 3. The media was replaced with half-fresh half-conditioned media after 3 h and the cells were used 4 days later (DIV 7). To validate protein knockdown (MOI 7.5), neurons were lysed in lysis buffer (20 mM HEPES pH 7.4, 100 mM NaCl, 1% Triton X-100, 5 mM EDTA, HALT Protease Inhibitor), resolved by SDS–PAGE and processed for Western blotting using anti-VAMP7 mouse monoclonal and anti-β-actin mouse monoclonal as a loading control or anti-Syntaxin 4 rabbit monoclonal and anti-β-Tubulin 3 mouse monoclonal as a loading control.

### Immunostaining protocol

Cells were fixed in 4% paraformaldehyde (PFA) for 10 min at room temperature, washed twice in phosphate-buffered saline (PBS) with 0.1% TX100 and blocked with PBS containing 2% bovine serum albumin and 0.2% TX100 for 1 h at room temperature. Cells were incubated with primary antibody diluted in blocking buffer for 1 h at room temperature or overnight at 4°C. Cells were washed three times in PBS with 0.1% TX100 and incubated with secondary antibody diluted in blocking buffer for 1 h at room temperature. After washing, the cells were incubated with DAPI diluted in PBS for 10 min and mounted using DAKO fluorescence mounting media. For quantification, 10-20 images per coverslip were averaged.

### Confocal microscopy

Imaging was performed using the following microscopes: (1) Leica TCS SP8 confocal equipped with a plan-apochromat 100x oil objective (Lecia, NA = 1.4) and LAS X software. (2) Leica Stellaris 5 confocal equipped with a plan-apochromat 63x oil objective (Lecia, NA = 1.4) and LAS X software. (3) LSM710 equipped with a plan-apochromat 63x oil objective (Zeiss, NA = 1.4) and ZEN software (Zeiss). (4) LSM900 with Airyscan 2 equipped with a plan-apochromat 63x oil objective (Zeiss, NA = 1.4) and 20x objective (Zeiss, NA = 0.8) and ZEN software (Zeiss).

### Live cell imaging

For total internal reflection fluorescence (TIRF) microscopy experiments, neurons were loaded with 2 µM BODIPY 558/568 C12 (Red-C12) overnight at 37°C in a 5% CO2 incubator, washed twice in warm PBS and imaged live in Tyrode’s solution (124 mM NaCl, 3 mM KCl, 2 mM CaCl2, 1 mM MgCl2, 10 mM HEPES pH 7.3 and 5-mM D-glucose) or Hanks’ Balanced Salt Solution (HBSS) with 1 μM MK6-83 or 20 μM BAPTA-AM. Syntaxin 4 knockdown neurons used were transduced at an MOI 7.5. Neurons were imaged using the following microscopes: (1) TIRF Olym6us IX73 microscope equipped with a plan-apochromat 100x/1.49 NA (Olympus Canada), incubation system to maintain 37°C, humidity, and CO_2_ (Live Cell Instruments Inc.), Hamamastu EMCCD-9100 camera (Hamamatsu Photonics) and images were acquired with Volocity imaging software (Quorum Technologies, Canada). Timelapse movies were acquired at 12-21 frames/sec and displayed as a maximum intensity projection or as a kymograph. (2) Elyra 7 microscope equipped with an alpha plan-apochromat 63x/1.46 NA oil objective, incubation system to maintain 37°C, humidity and CO_2,_ and ZEN software (Zeiss). Timelapse movies were acquired at 67 frames/sec. To quantify the number of fusion events, 30 sec movies were compressed into a maximum intensity projection. Regions outside the soma were removed and background was subtracted using a rolling ball radius of 50 pixels. The images were thresholded using the Minimum algorithm and the number of Red-C12-positive puncta (putative fusion events) and total soma area was analyzed using the Fiji software. For each independent experiment, the number of Red-C12 puncta detected for each cell were normalized to soma area and then multiplied by the average soma area to obtain fusion events/soma. For LysoTracker co-localization experiments, neurons were loaded with 50 nM LysoTracker Blue DND-22 and 1 µM BODIPY 581/591 C11 (BD-C11) in HBSS for 30 min at 37°C in a 5% CO_2_ incubator, washed twice in warm PBS and imaged live in HBSS. Cells were imaged at 37°C in a 5% CO_2_ chamber using LSM900 with Airyscan multiplex processing.

### Co-localization Analysis

For PLIN2 co-localization, Neurons were incubated in complete media containing 2 ∝M Red-C12 for 1 h, washed twice in warm PBS and incubated with HBSS for 1 h. Cells were washed twice in PBS, fixed in 4% PFA and immuno-stained for PLIN2 as described above. Quantification was performed by creating binary masks of PLIN2 and Red-C12 channels and generating an overlap image using the ‘AND’ function in Fiji-Image. Percent co-localization was expressed as the number of overlapping puncta divided by the number of Red-C12 puncta. For Lysotracker blue co-localization, cells were treated and imaged live as described above. Quantification of co-localization between Lysotracker Blue and oxidized BODIPY-C11 (green emission) was performed as described above and expressed at the number of overlapping puncta divided by the number of BODIPy-C11. Soma and neurite were analyzed separately.

### Fatty acid transfer assays

Neurons were incubated with complete media containing 2 ∝M Red-C12 for 16 h, washed twice in warm PBS and incubated with fresh complete media for 1 h. Red-C12 labelled neurons and unlabelled glia on separate coverslips were washed twice with warm PBS and the coverslips were sandwiched together (facing each other) separated by paraffin wax and incubated in HBSS for 4 h at 37°C with or without DMSO or 1 ∝M MK6-83. VAMP7 and syntaxin 4 knockdown neurons used were transduced at an MOI 2.5 and 7.5, respectively. Images for quantification were taken using LSM710 (Zeiss). Images for figures were taken using the LSM900 (Zeiss). For quantification, maximum intensity projections of three-dimensional image stacks were generated. The images were thresholded using the Yen algorithm and the number of Red-C12-positive particles was analyzed using Fiji software. Red-C12 puncta was determined to be within the cell by increasing the brightness/contrast such that the contours of the cell become visible (Fig, S1 E). The contribution of Red-C12 leaching from the coverslip and/or wax has been previously found to be negligible in this assay (Ioannou, Jackson, et al., 2019).

### Fatty acid accumulation assays

Neurons were incubated with complete media containing 4 μM Red-C12 for 30 min at 37°C, washed three times in warm PBS and incubated with fresh neuron media for 30 min at 37°C. Neurons were washed twice in warm PBS and treated with or without DMSO, ddH_2_O, 1 μM MK6-83, 20 μM BAPTA-AM, 250 nM Bafilomycin A1 (BAF A1), 250 nM Bafilomycin B1 (BAF B1), 50 μM Chloroquine (CQ) or 10 mM 3-Methyladenine (3-MA) in HBSS for 30 min at 37°C. Neurons were fixed and imaged using LSM710 or LSM900 (Zeiss). VAMP7 and syntaxin 4 knockdown neurons used were transduced at an MOI 5 and only GFP-positive neurons were quantified. Quantification was the same as transfer assay.

### Fatty acid release assays

Neurons were grown on PDL-coated 6-well plastic dishes. Neurons were loaded with or without 2 μM Red-C12, 5 μg/ml CholEsteryl BODIPY 542/563 C11 (BD-CE) or 5 μg/ml BODIPY 493/503 (BD493) for 16 h in complete media. Cells were washed three times in warm PBS, incubated with fresh media for 1 h and treated with or without DMSO, ddH_2_O, 1 μM MK6-83, 20 μM BAPTA-AM, 50 μM α-tocopherol, 10 μM Ferrostatin-1, 1 μM Rotenone, 100 μM Erastin, 100 μM Kainate, 500 μM NMDA, 250 nM Bafilomycin A1 (BAF A1), 250 nM Bafilomycin B1 (BAF B1), 50 μM Chloroquine (CQ) or 10 mM 3-Methyladenine (3-MA) in HBSS or complete medium prepared with phenol-red free Neurobasal medium for 4 h at 37°C. Neuron-conditioned media was collected and centrifuged at 16,000 x g for 15 min to remove dead cells. The supernatant was analyzed for Red-C12, BD-CE or BD493 fluorescence using Synergy Mx Multi-Mode Microplate Reader (BioTek Instruments Inc.). VAMP7 and syntaxin 4 knockdown neurons used were transduced at an MOI 7.5. Each well was analyzed in triplicate and averaged for quantification.

### Lipid droplet analysis in *Drosophila* retinas

Expression of *UAS-RNAi* was induced using *da-GAL4* (Wodarz et al., 1995), *Rh1-GAL4* (Mollereau et al., 2000), or *54C-GAL4* (Nagaraj & Banerjee, 2007). *UAS-RNAi* lines (stock numbers *luciferase, 31603*; *Vamp7*, 38300 and 43543; *Syx4*, 31667 and 31198) were obtained from the Bloomington *Drosophila* Stock Center (BDSC). *Rh:ND42 RNAi* was generated previously (Liu et al., 2015) and was made available from BDSC under the stock number 76598. Whole-mount staining of fly retinas with Nile Red to visualize lipids was performed as in Moulton et al., (2021). In brief, retinas were isolated after fixation in 3.7% formaldehyde overnight, rinsed with PBS, and stained for 20 min in 1 mg/mL Nile Red in PBS. After staining, retinas were rinsed with PBS, mounted in Vectashield (Vector Labs) on slides, and imaged on a Leica SP8 confocal microscope using a 63x glycerol submersion lens with 3x zoom. ImageJ was utilized to visualize fly retinal images and all genotypes were blinded prior to quantification. Lipid droplets with diameter ≥0.5 µM were manually quantified.

### Quantitative real-time PCR (qPCR) on *Drosophila*

qPCR was performed as previously described (Goodman et al., 2019). Briefly, total RNA was extracted from larvae or adult heads using Trizol. Samples were DNAse treated using the TURBO DNA-free Kit (ThermoFisher). cDNA was synthesized using the High-Capacity cDNA Reverse Transcription Kit (ThermoFisher). Mean fold change was determined using the ΔΔCt method from data collected using an Applied Biosystems’s QuantStudio 5 Real-Time PCR system. All primers used are listed in the key resource table and are intron spanning with 90-110% efficiency by serial dilution curve.

### ROS and lipid peroxidation assays

For CellRox assay, neurons were washed three times in warm PBS and incubated in either complete media or HBSS for 4 h with 5 ∝M CellROX Green added for the last 30 min. Cells were fixed and imaged using Leica SP8. Mean intensities for thresholded nuclei were measured using Fiji-ImageJ. For BD-C11 lipid peroxidation assay, neurons were washed three times in warm PBS and incubated in complete media or HBSS with or without DMSO, 1 ∝M MK6-83, 50 ∝M α-tocopherol or 10 ∝M Ferrostain-1 for 4 h with 2 ∝M BD-C11 added for the last 60 min. Cells were fixed and imaged using the Zeiss LSM710. Images for figures were taken using the Zeiss LSM900. Relative lipid peroxidation was indicated by the ratio of green fluorescence intensity over red fluorescence intensity using Fiji-ImageJ. For lipid hydroperoxide assay, neurons were grown on PDL-coated 6-well plastic dishes. Neurons were washed three times in warm PBS and incubated in HBSS with or without DMSO, 1 ∝M MK6-83, 20 ∝M BAPTA-AM, 10 ∝M Ferrostain-1, 250 nM Bafilomycin A1 (BAF A1) or 10 mM 3-Methyladenine (3-MA) for 6 h. Lipid hydroperoxides were extracted from neuronal lysates and neuron-conditioned media and analyzed using a lipid hydroperoxide assay kit according to the manufacturers protocol and Synergy Mx Multi-Mode Microplate Reader (BioTek). VAMP7 and syntaxin 4 knockdown neurons used for the lipid hydroperoxide assay were transduced at an MOI 7.5.

### Iron assays and lysosome function assays

Neurons were grown on PDL-coated 6-well plastic dishes. Neurons were washed twice in warm PBS and incubated in HBSS with or without ddH_2_O or 10 mM 3-Methyladenine (3-MA) for 6 h. VAMP7 and syntaxin 4 knockdown neurons used were transduced at an MOI 7.5. Total iron (Fe^2+^ and Fe^3+^) from neuronal lysates and neuron-conditioned media was measured using an iron assay kit according to the manufacturers protocol. Neuronal lysates were analyzed using a cathepsin D activity assay kit according to the manufacturers protocol. Assay detection was performed using the Synergy Mx Multi-Mode Microplate Reader (BioTek).

### Fatty acid, lipid hydroperoxide and iron depletion assays

Neurons were plated on 10-cm PDL-coated plastic plates. For the fatty acid depletion assay, neurons were incubated with complete media containing 2 ∝M Red-C12 for 16 h. Neurons were washed twice in warm PBS, incubated for an additional 1 h in complete media followed by 12 h in HBSS. Neuron-conditioned media was centrifuged at 10,000 x g for 20 min (low g) at 4°C to remove dead cells or cell debris. The supernatant was centrifuged at 200,000 x g for 2.5 h (high g) at 4°C in a TLA-100.3 fixed angle rotor (k-factor 14) (Beckman Coulter) and Red-C12 fluorescence of the starting material, low g supernatant and high g supernatant was measured using Synergy Mx Multi-Mode Microplate Reader (BioTek Instruments Inc.). For the lipid hydroperoxide depletion assay, neurons were washed twice in warm PBS and incubated in 5 mL HBSS with or without 1 ∝M MK6-83 for 6 h. Neuron-conditioned media was centrifuged at 10,000 x g for 20 min (low g) at 4°C and the pellet was resuspended in 200 ∝l of HBSS. The supernatant was centrifuged at 200,000 x g for 2.5 h (high g) at 4°C and the pellet was resuspended in 200 ∝l HBSS. Lipid hydroperoxides were extracted from the starting material, low g pellet, high g pellet and high g supernatant using a lipid hydroperoxide assay kit and Synergy Mx Multi-Mode Microplate Reader (BioTek Instruments Inc.). For the iron depletion assay, neurons were washed twice in warm PBS and incubated in 5 mL HBSS with or without ddH_2_O or 10 mM 3-Methyladenine (3-MA) for 6 h. Neuron-conditioned media was centrifuged at 10,000 x g for 20 min (low g) at 4°C and the pellet was resuspended in 200 ∝l of HBSS. The supernatant was centrifuged at 200,000 x g for 2.5 h (high g) at 4°C and the pellet was resuspended in 200 ∝l HBSS. Total iron (Fe^2+^ and Fe^3+^) of the starting material, low g pellet, high g pellet and high g supernatant was measured using an iron assay kit and Synergy Mx Multi-Mode Microplate Reader (BioTek Instruments Inc.).

### Total cell death and TUNEL apoptosis assays

For total cell death assay, neurons were plated on 96-well PDL-coated plastic plates. Neurons were washed twice in warm PBS and treated with or without 1 μM MK6-83, 20 μM BAPTA-AM, 50 μM α-tocopherol, 10 μM Ferrostatin-1, 10 μM Erastin (HBSS) or 100 μM Erastin (complete media) in HBSS or phenol-red free Neurobasal medium (complete media) for 12 h. Neurons were washed twice in warm PBS and incubated with 4 μM Ethidium Homodimer-1 in PBS for 30 min at 37°C to label dead cells, washed twice in PBS and fluorescence was measured using Synergy Mx Multi-Mode Microplate Reader. VAMP7 and syntaxin 4 knockdown neurons used were transduced at an MOI 7.5. For terminal deoxynucleotidyl transferase dUTP nick end labelling (TUNEL) apoptosis assay, neurons were washed twice in warm PBS and treated in complete media or HBSS with or without DMSO, 1 ∝M MK6-83 or 50 ∝M α-tocopherol for 12 h. Cells were washed twice, fixed in 4% PFA, stained for apoptotic cells using a TUNEL assay kit according to the manufacturers protocol (Abcam; ab66110) and imaged within 3 h of staining using Zeiss LSM900. VAMP7 and syntaxin 4 knockdown neurons used were transduced at an MOI 7.5 For quantification, maximum intensity projections of three-dimensional image stacks were generated. BrdU-positive apoptotic cells versus total cells were detected by thresholding images using the Yen algorithm and the number of particles was analyzed using Fiji software. For quantification, 5 images per coverslip were averaged.

### Autophagy LC3 assay

Neurons were grown on PDL-coated 6-well plastic dishes. Neurons were washed twice in warm PBS and treated with or without 250 nM Bafilomycin A1 (BAF A1), 250 nM Bafilomycin B1 (BAF B1), 50 μM Chloroquine (CQ), 10 mM 3-Methyladenine (3-MA) or 10 μM Erastin in HBSS for 6 h. Neurons were washed twice in PBS and lysed in lysis buffer, resolved by SDS–PAGE and processed for Western blotting using anti-LC3B rabbit polyclonal and anti-β-actin mouse monoclonal as a loading control.

### 3-MA extracellular vesicle release assay

Neurons were plated on 10-cm PDL-coated plastic plates. Neurons were washed twice in warm PBS and treated with or without ddH_2_O or 10 mM 3-Methyladenine (3-MA) for 12 h in HBSS. Neurons were washed twice in PBS and lysed in lysis buffer. Neuron-conditioned media was centrifuged at 10,000 x g for 20 min (low g) at 4°C and the pellet was resuspended in 25 ∝l lysis buffer. The supernatant was centrifuged at 200,000 x g for 2.5 h (high g) at 4°C in a TLA-100.3 fixed angle rotor (k-factor 14) (Beckman Coulter) and the pellet was resuspended in 25 ∝l lysis buffer. Samples were resolved by SDS–PAGE and processed for Western blotting using anti-syntenin rabbit polyclonal.

### Transmission electron microscopy

For in-situ studies of hippocampal neurons, 2-month-old male C57/BL6N mice were anesthetized and perfused with 3% PFA (60 mM NaCl, 130 mM glycerol, 10 mM sodium phosphate buffer). The brain was dissected and postfixed with 3% PFA (30 mM of NaCl, 70 mM glycerol, 30 mM PIPES buffer, 10 mM betaine, 2 mM CaCl2, 2 mM MgSO4) under RT for 2 h. The brain was rinsed in a 400 mOsM buffer (65 mM NaCl, 100 mM glycerol, 30 mM PIPES buffer, 10 mM betaine, 2 mM CaCl2, and 2 mM MgSO4) for 30 min, and sectioned by vibratome (coronal sections, 100 mm) using a Leica VT1000S vibratome in the same buffer. Sections were fixed in 1% PFA, 2% glutaraldehyde solution (30 mM NaCl, 70 mM glycerol, 30 mM PIPES buffer, 10 mM betaine, 2 mM CaCl2, 2 mM MgSO4, 75 mM sucrose) overnight at 4°C. Sections were washed in 400 mOsM buffer. Round samples of the hippocampus were created using a 2 mm biopsy punch (Miltex). The 2 mm samples were dipped in hexadecene, placed in 100 μm aluminum carrier, covered with a flat carrier and high-pressure frozen using a Wohlwend compact 01 High pressure freezer. Samples were then freeze-substituted in 0.5% osmium tetroxide, 20 mM 3-Amino-1,2,4-triazole, 0.1% uranyl acetate, 4% water in acetone, using a Leica AFS2 system. Specimens were further dehydrated in 100% acetone and embedded in Durcupan resin. Ultrathin sections (60 nm) were imaged in a Tecnai Spirit electron microscope (FEI, Hillsboro, OR) operating at 80 kV equipped with an Ultrascan 4000 digital camera (Gatan Inc, CA).

For in vitro studies, primary hippocampal neurons were plated on 2 x 2 PDL-coated aclar sheets. Neurons were washed twice in warm PBS and fixed in 1% PFA, 2% glutaraldehyde in 0.1M CaB (0.1-0.2 M Cacodylate buffer pH 7.4) containing 4% polyvinylpyrrolidone and 0.05% CaCl_2_ for 16 h at 4°C. Cells were washed with 0.1 M CaB and post-fixed with 2% osmium-imidazole buffer, pH 7.5. Samples were dehydrated in a graded series of ethanol, 30%, 50% and then block stained with 1% uranyl acetate in 70% ethanol overnight. Samples were then further dehydrated in 70%, 95% and 100% ethanol and anhydrous acetone and embedded in Durcupan (ACM) resin. Blocks were sectioned using an ultramicrotome (Leica, EM UC6; 70 nm thickness) and placed on a 400 mesh copper grid. Sections were stained with 1% uranyl acetate and 1% lead citrate and imaged using a transmission electron microscope (JEOL JEM-2100, Gatan Orius camera with Digital micrograph) at 200kV acceleration voltage.

### Cryo-electron microscopy

Neurons were plated on 10-cm PDL-coated plastic plates. Neuron-conditioned medium was centrifuged at 10,000×g at 4°C for 20 min. The supernatant was centrifuged at 300,000×g at 4°C for 3 h in a TLA-110 fixed angle rotor (k-factor 13) (Beckman Coulter). The pellet was resuspended in 10 µL PBS of which 3 µL was adsorbed onto glow-discharged 2 nm carbon layer coated Quantifoil R1.2/1.3 400 mesh gold grids, blotted for 3 s at 4°C and targeted relative humidity level of 100% and plunge frozen using a Vitrobot Mark IV (Thermo Fisher Scientific, MA) or for 4–6 s at 4°C and ~ 86% and plunge frozen using a Leica EM GP (Leica Microsystems Inc, Buffalo Grove, IL). The samples were imaged in an FEI Titan Krios electron microscope operated at 300 kV with a K2 Summit camera and energy filter (Gatan, Inc., CA). Images were recorded at a pixel size of 2.208 Å/px (1.104 Å/px super-resolution) at a dose rate of 10 e^-^/px/s over 9.9 s using 0.3 s frames (20.7 e^-^/Å^2^ total dose, 0.62 e^-^/Å^2^ per frame) using SerialEM (Mastronarde, 2005). Movies were binned to a pixel size of 2.208 Å/px and aligned using CisTEM (Egelman et al., 2018). The diameter of 125 particles was quantified manually using Fiji-imageJ.

### Quantification and statistical analyses

Datasets were assembled in Microsoft Excel 365 (Microsoft Corp.). Statistical analysis and graphing were performed using Graphpad Prism 9. All graphs are depicted as Superplots where the biological replicates are shown in large shapes, and the corresponding technical replicants shown as small shapes of the same color (Lord et al., 2020). Statistical test used and p-values can be found in the figures and/or figure legends. Where stated the Bonferroni correction method was used to correct for multiple comparisons (raw p-values were multiplied by the number of comparisons in the experiment).

---

**Figure S1.**
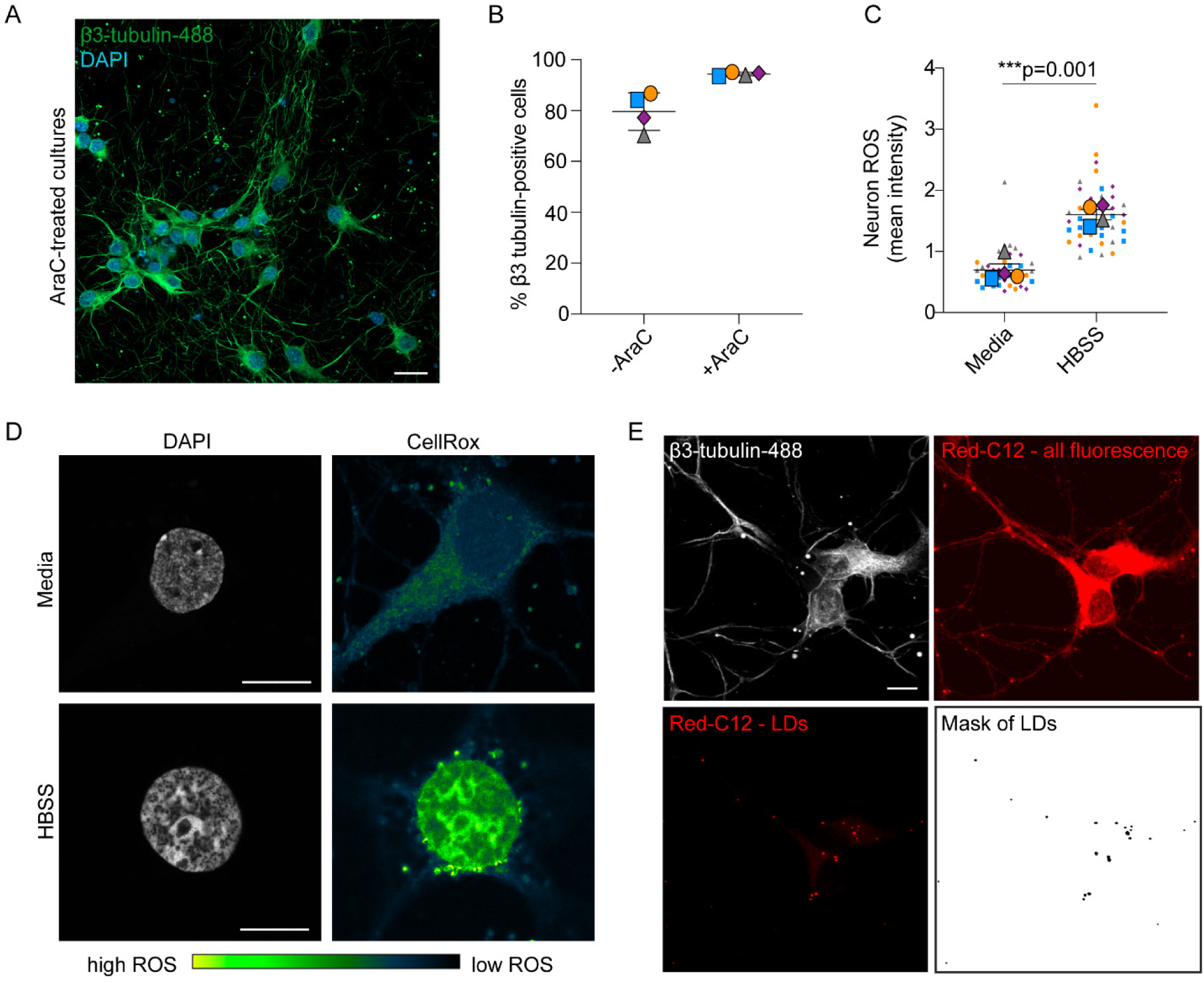
Neurons increase oxidative stress in HBSS. (A) Neuronal cultures grown in AraC (cytosine arabinoside) were fixed and immunostained for neuron-specific β3-tubulin. Scale bar, 20 ∝m. (B) Percentage of β3-tubulin-positive neurons. n=4 independent experiments; mean ± SD. (C and D) Confocal image of neurons in media or HBSS labelled with CellRox Green. Mean intensity of nuclear CellRox Green was quantified. n=4 independent experiments; mean ± SEM. Student’s T-test. (E) Representative image showing how saturated Red-C12 display labels the whole cell including neurites and allows for the quantification of Red-C12 puncta specifically within cells.

**Figure S2.**
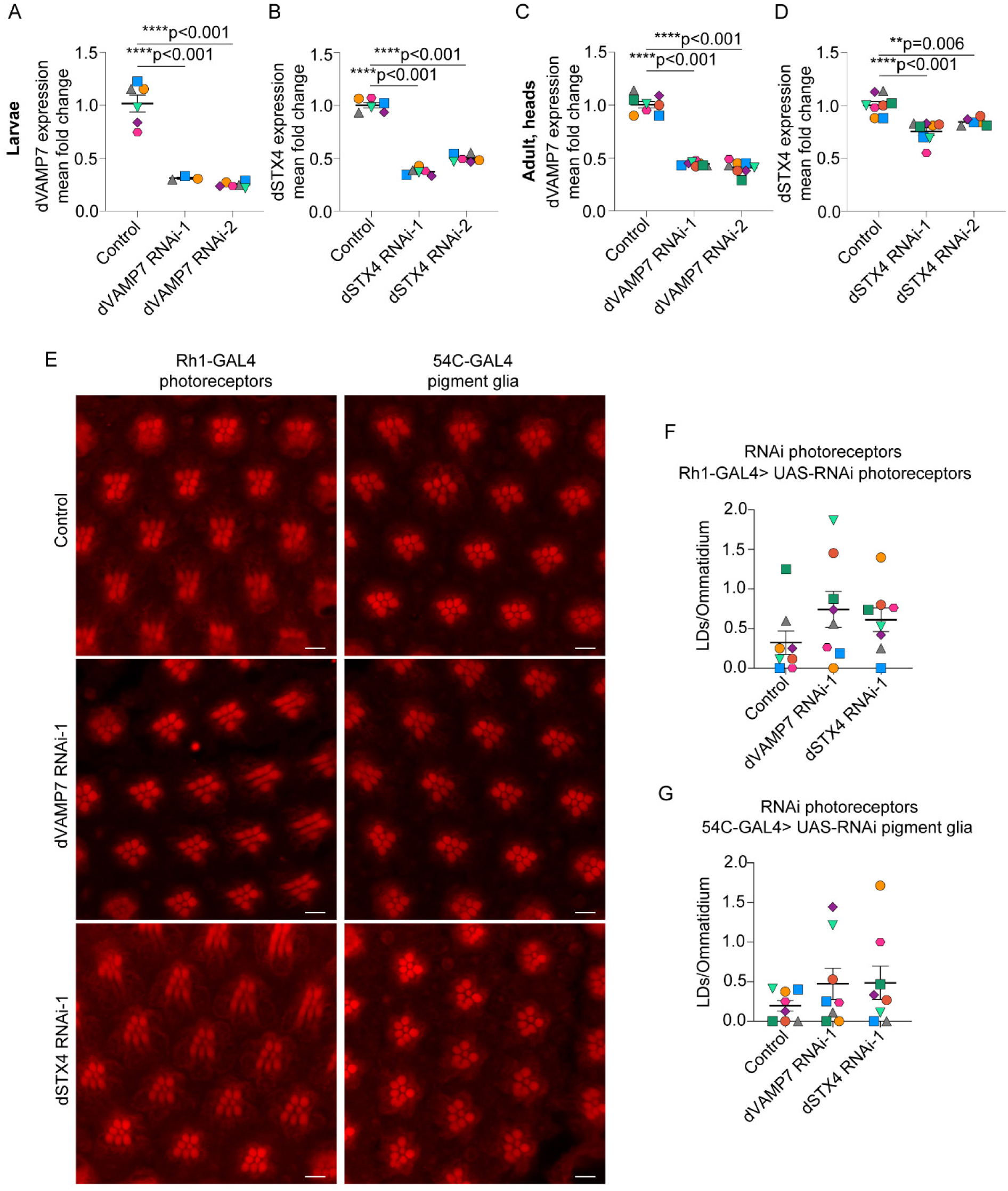
VAMP7 or Syntaxin 4 knockdown in the absence of ROS does not induce glial lipid droplets *in vivo.* (A – D) *UAS-RNAi* directed against d*VAMP7*, d*STX4 or* d*LUC* (control) genes was induced using the ubiquitous driver, *Da-GAL4*, and total RNA from either L3 larvae or 1-2d adult heads was quantified by qPCR. Minimum n = 3; mean ± SEM. One-way ANOVA with Dunnett’s post test. (E) Confocal image. No Nile Red positive lipid droplets (LD) in pigment glial cells in the absence of ROS in photoreceptors. *UAS-RNAi* directed against d*VAMP7* and d*STX4* genes was induced in neurons using *Rh1-GAL4* and in pigment glia using *54C-GAL4*. Control fly is UAS-empty. Scale bars, 5 ∝m. (F) Number of LDs per ommatidium when d*VAMP7* and d*STX4* is knocked down in photoreceptor neurons (Rh1-GAL4). n = 8 animals per genotype; mean ± SEM. One-way ANOVA with Tukey’s post test. (G) Number of lipid droplets per ommatidium when d*VAMP7* and d*STX4* is knocked down in pigment glia (*54C-GAL4*). n = 8 animals per genotype; mean ± SEM. One-way ANOVA with Tukey’s post test.

**Figure S3.**
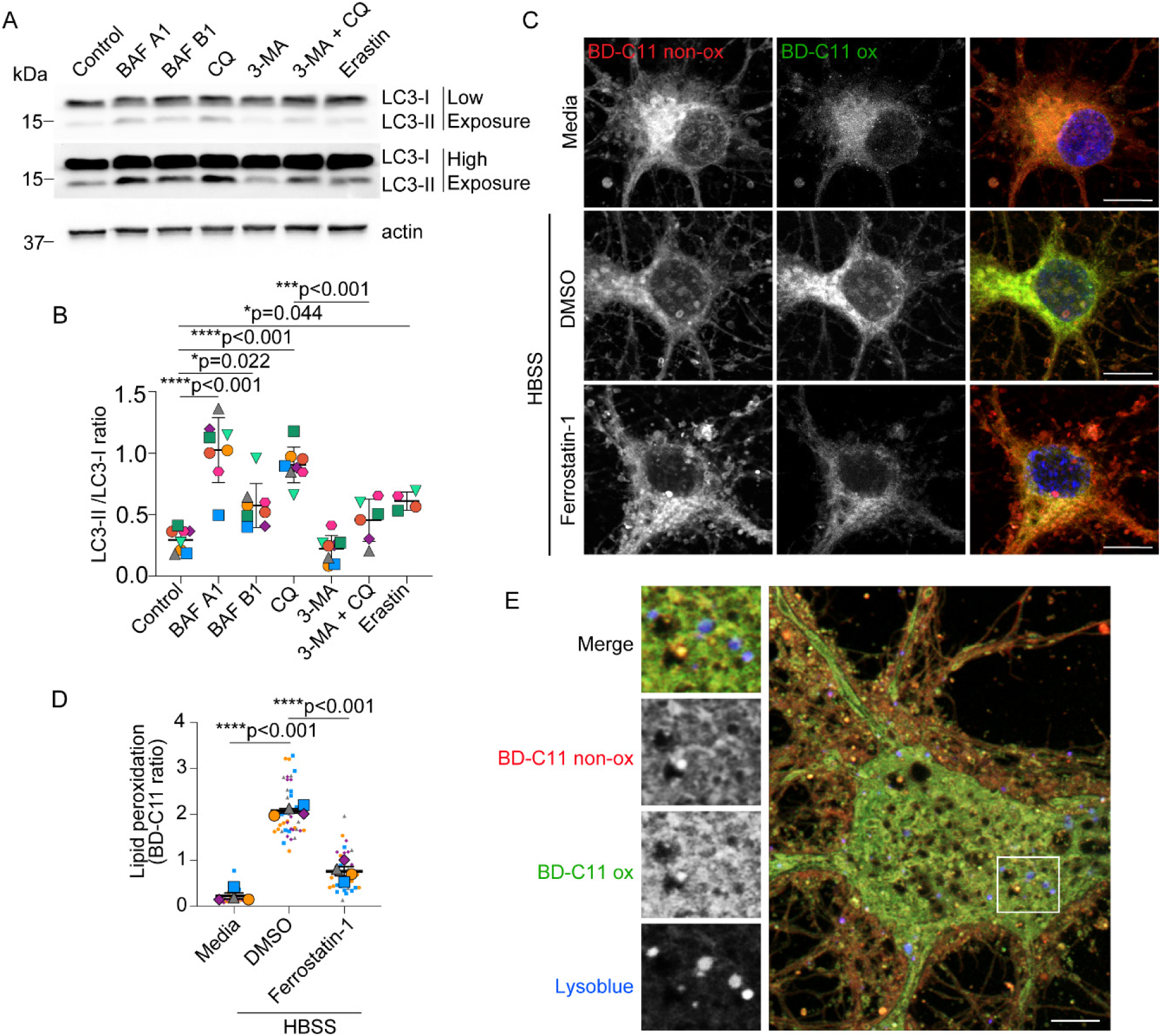
Validation of autophagy targeting drugs in neurons and increased peroxidated lipids in neurons treated with HBSS. (A and B) Neurons ± bafilomycin A1 (BAF A1), bafilomycin B1 (BAF B1), chloroquine (CQ), 3-Methyladenine (3-MA) or erastin in HBSS were analyzed by Western blot for LC3 and β-actin. n = 4-6 independent experiments; mean ± SD. One-way ANOVA with Tukey’s post test. (C) Confocal maximum intensity projection of neurons in media or HBSS ± DMSO or ferrostatin-1 labelled with BD-C11. Total (BD-C11 non-ox) and peroxidated lipids (BD-C11-ox). Scale bars, 10 ∝m. (D) Quantification of lipid peroxidation (ratio BD-C11 ox/ BD-C11 non-ox) in neurons in media or HBSS ± DMSO or ferrostatin-1. n = 4 independent experiments; mean ± SEM. One-way ANOVA with Tukey’s post test. (E) Airyscan image of live neuron in HBSS labelled with BD-C11 and LysoTracker Blue. Total (BD-C11 non-ox) and peroxidated (BD-C11 ox) lipids. Boxed area magnified in left panels. Scale bar, 5 ∝m.

**Figure S4.**
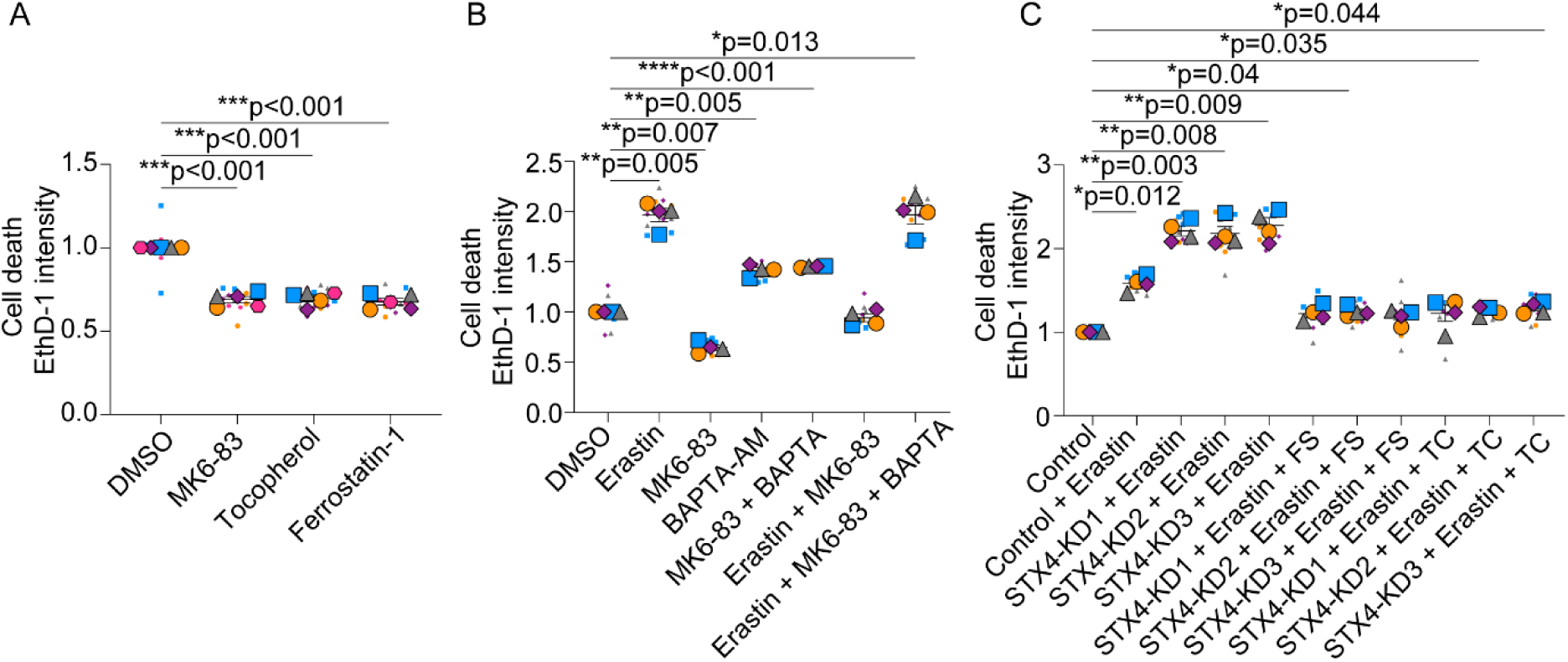
Autolysosomal exocytosis reduces neuronal death. (A) Neurons were treated with HBSS ± DMSO, MK6-83, α-tocopherol or ferrostatin-1 and stained with ethidium homodimer-1 (EthD-1). n = 5 independent experiments; mean ± SEM. One sample t-test with Bonferroni correction. (B) Neurons were treated with HBSS ± DMSO, erastin, MK6-83 or BAPTA-AM and stained with EthD-1. n = 4 independent experiments; mean ± SEM. One sample t-test with Bonferroni correction. (C) Neurons were transduced with lentivirus expressing non-targeting shRNAmiR (control) or shRNAmiRs targeting syntaxin 4 (STX4). Neurons were treated with media ± DMSO, erastin, ferrostatin-1 or α-tocopherol and stained with EthD-1 and normalized to non-targeting control neurons. n = 4 independent experiments; mean ± SEM. One sample t-test with Bonferroni correction.

**Figure S5.**
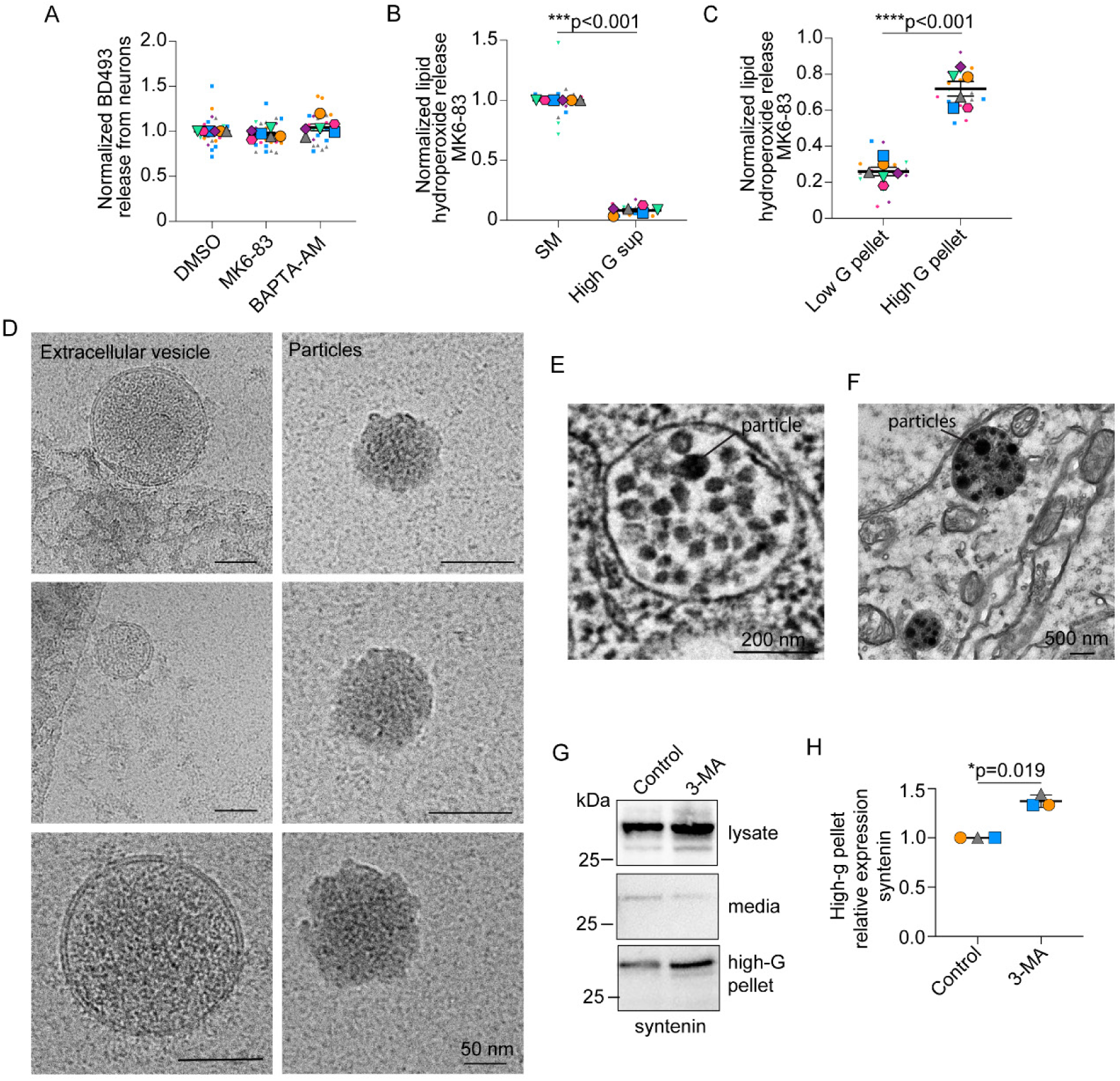
Peroxidated lipids released as lipid-protein particles. (A) Neuron-conditioned HBSS analyzed for BD493 fluorescence and normalized to DMSO treated control neurons. n = 6 independent experiments; mean ± SEM. One sample t-test with Bonferroni correction. (B) Neuron-conditioned HBSS analyzed for lipid hydroperoxides following MK6-83 treatment and centrifugation. SM, starting material. n = 6 independent experiments; mean ± SEM. One sample t-test with Bonferroni correction. (C) Pellets obtained from centrifugation of MK6-83 treated neuron-conditioned HBSS were analyzed for lipid hydroperoxides. n = 6 independent experiments; mean ± SEM. Student’s t-test. (D) Pellets obtained from high G centrifugation of neuron-conditioned media were examined by cryoEM. Scale bars, 50 nm. (E) Cultured neurons stained with imidazole-buffered osmium were imaged by transmission electron microscopy. Scale bar, 200 nm (F) Hippocampal neuron from sectioned mouse brain, stained with imidazole-buffered osmium were imaged by transmission electron microscopy. Scale bar, 500 nm. (G and H) Neuronal lysates and neuron-conditioned HBSS ± 3-Methyladenine (3-MA) were analyzed by Western blot for syntenin levels following centrifugation. SM, starting material. n = 3 independent experiments; mean ± SD. One sample t-test adjusted with Bonferroni correction.

